# How to build a water-splitting machine: structural insights into photosystem II assembly

**DOI:** 10.1101/2020.09.14.294884

**Authors:** Jure Zabret, Stefan Bohn, Sandra K. Schuller, Oliver Arnolds, Madeline Möller, Jakob Meier-Credo, Pasqual Liauw, Aaron Chan, Emad Tajkhorshid, Julian D. Langer, Raphael Stoll, Anja Krieger-Liszkay, Benjamin D. Engel, Till Rudack, Jan M. Schuller, Marc M. Nowaczyk

## Abstract

Biogenesis of photosystem II (PSII), nature’s water splitting catalyst, is assisted by auxiliary proteins that form transient complexes with PSII components to facilitate stepwise assembly events. Using cryo-electron microscopy, we solved the structure of such a PSII assembly intermediate with 2.94 Å resolution. It contains three assembly factors (Psb27, Psb28, Psb34) and provides detailed insights into their molecular function. Binding of Psb28 induces large conformational changes at the PSII acceptor side, which distort the binding pocket of the mobile quinone (Q_B_) and replace bicarbonate with glutamate as a ligand of the non-heme iron, a structural motif found in reaction centers of non-oxygenic photosynthetic bacteria. These results reveal novel mechanisms that protect PSII from damage during biogenesis until water splitting is activated. Our structure further demonstrates how the PSII active site is prepared for the incorporation of the Mn_4_CaO_5_ cluster, which performs the unique water splitting reaction.

**One Sentence Highlight:** The high-resolution Cryo-EM structure of the photosystem II assembly intermediate PSII-I reveals how nature’s water splitting catalyst is assembled, protected and prepared for photoactivation by help of the three assembly factors Psb27, Psb28 and Psb34.

## Introduction

Photosystem II (PSII) is the only enzyme that catalyzes the light-driven oxidation of water, a thermodynamically demanding reaction that drives photosynthesis, sustaining life on our planet (Hohmann-Marriott and Blankenship, 2011; Sanchez-Baracaldo and Cardona, 2020; Vinyard et al., 2013). This multi-subunit membrane protein complex is located in the thylakoid membranes of cyanobacteria, algae and plants. PSII strips electrons from water and injects them into the photosynthetic electron transport chain (PET), where they are transferred via plastoquinone towards cytochrome b_6_f and further via plastocyanin to photosystem I (PSI). PSI performs another light-driven charge separation, which enables the reduction of ferredoxin and the synthesis of NADPH—a universal redox mediator. At the same time, the PET drives unidirectional transport of protons through the thylakoid membrane, generating an electrochemical gradient used for ATP synthesis. Photosynthetic organisms use this light-generated ATP and NADPH to perform carbon fixation via the Calvin-Benson cycle, which is the main biochemical pathway for the conversion of atmospheric CO_2_ into organic compounds. This photosynthetic pathway, and the web of life it sustains, all begins with PSII using the energy of sunlight to split water.

PSII forms a homodimer with a molecular mass of ∼500 kDa (Boekema et al., 1995), with each monomer composed of at least 20 protein subunits and numerous cofactors, including chlorophylls, quinones, carotenoids, lipids, bicarbonate and the unique Mn_4_CaO_5_ cluster that splits water into oxygen and protons (Cox et al., 2020; Shen, 2015; Yano et al., 2015). The two core proteins D1 and D2 form a central, membrane-intrinsic heterodimer, which binds all important redox cofactors involved in internal electron transfer (Ferreira et al., 2004). Light-excitation leads to a charge-separated state in which an electron is transferred from the chlorophyll assembly P_680_ (Cardona et al., 2012) to the nearby pheophytin (Holzwarth et al., 2006). Subsequently, the electron is passed to the bound plastoquinone (Q_A_) and then to the mobile plastoquinone molecule (Q_B_), which leaves the complex after accepting two electrons and two protons (Müh et al., 2012). The electron hole at P_680_ is filled by oxidation of an adjacent tyrosine residue (Tyr_Z_) (Faller et al., 2001) and finally by the oxygen evolving complex (OEC) that contains the Mn_4_CaO_5_ cluster. In cyanobacteria, the cluster is shielded at the luminal side by the three extrinsic proteins, PsbO, PsbU and PsbV, which regulate access to the OEC by forming a complex network of channels for different substrates and products (Roose et al., 2016). Light energy is collected and funneled towards P_680_ by the two membrane-intrinsic antenna proteins CP43 and CP47. These proteins bind most of the chlorophyll molecules and are located at opposite sides of the D1/D2 heterodimer (Müh and Zouni, 2020). Moreover, at least twelve small transmembrane subunits with one or two transmembrane helices have been identified in PSII (Shi et al., 2012), including cytochrome-b_559_ (Stewart and Brudvig, 1998).

Structural and spectroscopic investigations have revealed these aforementioned comprehensive insights into PSII function (Cox et al., 2014; Kern et al., 2018; Kupitz et al., 2014; Suga et al., 2019; Umena et al., 2011), but we are far from understanding PSII biogenesis with molecular detail. How nature facilitates the assembly of a multi-subunit, multi-cofactor membrane protein complex is a fundamental unsolved question. The biogenesis of PSII is even more challenging, as the mature complex performs sophisticated and extreme redox chemistry to catalyze the light-driven oxidation of water. This can easily lead to the formation of reactive oxygen species (e.g., singlet oxygen is produced by triplet chlorophyll in the PSII reaction center) and subsequent loss of function due to damaged proteins and cofactors (Krieger-Liszkay et al., 2008; Vass, 2012). Biogenesis intermediates with only partially functional fragments of the redox chain are particularly prone to damage, thus demanding specialized protection mechanisms for the assembly process. Therefore, PSII biogenesis is not a spontaneous process but rather must be tightly regulated by the action of assembly factors. Thus far, more than 20 auxiliary proteins have been identified that guide the stepwise assembly of PSII subunits and cofactors via intermediate modules, which are assembled independently and then joined together to produce mature PSII (Heinz et al., 2016; Nickelsen and Rengstl, 2013; Nixon et al., 2010). In cyanobacteria, PSII biogenesis begins with the formation of the D1/D2 heterodimer reaction center (RC) complex from the D1 precursor protein (pD1) and the D2 protein. This is assisted by the PSII assembly factor Ycf48 after partial processing of the D1 C-terminal extension by the D1 specific peptidase CtpA (Komenda et al., 2007; Komenda et al., 2008). In the next step, the assembly factor Psb28 helps CP47 join the RC complex to form the RC47 complex, in which iD1 is further processed to its mature form by CtpA (Boehm et al., 2012; Dobáková et al., 2009). Almost all ligands of the Mn_4_CaO_5_ cluster are already present at this stage, except for those provided by CP43, which comes pre-constructed with assembly factor Psb27 and several small subunits (together called the CP43 module) (Komenda et al., 2012). Psb28 is released as CP43 binds, and the resulting Psb27-PSII monomer is activated by maturation of the OEC and the binding of the extrinsic proteins PsbO, PsbU and PsbV (Mamedov et al., 2007; Nowaczyk et al., 2006; Roose and Pakrasi, 2008). Finally, PSII biogenesis completes with dimerization of two fully assembled monomers and attachment of the soluble phycobilisome antenna complexes. Interestingly, deletion of *psbJ*, which encodes a small single transmembrane helix protein at the entrance of the PSII plastoquinone channel, leads to massive accumulation of an intermediate monomeric PSII complex, which contains both assembly factors Psb27 and Psb28 (Nowaczyk et al., 2012).

Physiological studies of Psb27 and Psb28 deletion strains point towards multifaceted functions. Cyanobacterial mutants lacking Psb28 exhibited slower autotrophic growth, particularly under stress conditions (Dobáková et al., 2009; Sakata et al., 2013), and limited synthesis of Chl-binding proteins but without decrease in PSII functionality (Dobáková et al., 2009). The Psb28 mutant also exhibited an overall increase in PSII repair and faster recovery from photodamage (Dobáková et al., 2009). Chemical cross-linking combined with mass spectrometry revealed that Psb28 binds to the cytosolic side of CP47 close to cytochrome-b559 and the Q_B_ binding site. Based on this, researchers postulated a protective role for Psb28, whereby it blocks electron transport to the acceptor side of PSII, thereby protecting the RC47 complex from excess photodamage during the assembly process (Weisz et al., 2017). This hypothesis is strengthened by the observation that Psb28 is also found in PSII repair complexes (Bečková et al., 2017). The luminal PSII assembly factor Psb27 has been similarly well investigated. This lipoprotein is predominantly associated with inactive PSII fractions involved in assembly or repair (Bečková et al., 2017; Bentley et al., 2008; Grasse et al., 2011; Komenda et al., 2012; Liu et al., 2011b; Nowaczyk et al., 2006; Weisz et al., 2019), stabilizing the CP43 luminal domain and presumably facilitating the assembly of the OEC.

Our current knowledge of PSII biogenesis mainly describes the order of events and protein composition of each intermediate, as well as the general roles of PSII assembly factors. However, the precise molecular functions of these intermediate complexes and the involved assembly factors are still elusive due to their low abundance and intrinsic instability. High-resolution structural information is of vital importance to gain a deeper understanding into the molecular action of PSII assembly factors, as they are proposed to alter the structures of their associated PSII proteins to provide protection or facilitate specific biogenesis transitions.

Here, we use cryo-EM single particle analysis to describe the first molecular structure of a PSII assembly intermediate. This structure represents one of the key transitions in PSII biogenesis: the attachment of the CP43 module to the pre-assembled RC47 reaction center complex, which precedes incorporation and activation of the Mn_4_CaO_5_ cluster. We complement this structural data with spectroscopic analysis, revealing the first detailed insights into the molecular mechanisms of PSII assembly. Our study provides mechanistic answers to three long-standing questions: i) How do assembly factors modulate the structures of PSII subunits to assist biogenesis? ii) How is PSII protected from photodamage during assembly? iii) How is the PSII active site prepared for incorporation of the Mn_4_CaO_5_ cluster?

## Results

### Structure determination of the PSII assembly intermediate (PSI-I)

Stable PSII intermediates were purified from the *T. elongatus* Δ*psbJ* mutant (Nowaczyk et al., 2012) by affinity chromatography using a twin-strep-tag fused to the C-terminus of the CP43 subunit and subsequent ion exchange chromatography (Fig. S1A). The main peak of the IEC profile corresponds primarily to monomeric PSII, which lacks the extrinsic subunits PsbO, PsbU and PsbV that are indicative for water splitting activity (Fig. S1B and C). Single particle cryo-EM analysis of this PSII fraction resulted in three different high-resolution maps that allowed model building with high confidence and excellent statistics (Fig. S2, Table S1). In addition to the protein subunits, we also faithfully assigned all essential non-protein cofactors, including chlorophylls, quinones, carotenoids and lipids, which are also present in the mature PSII complex (Fig. S3). Consistent with previous biochemical studies (Grasse et al., 2011; Nowaczyk et al., 2006; Nowaczyk et al., 2012), the EM density corresponding to the fully assembled, active Mn_4_CaO_5_ cluster is missing in the purified biogenesis intermediates. The first cryo-EM map (2.94 Å), which we call PSII-I (for PSII-Intermediate), provides a snapshot of the attachment of the CP43 module to the pre-assembled RC47 reaction center complex (Fig. 1). This PSII intermediate contains three assembly factors (Psb27, Psb28 and Psb34), as well as almost all the membrane-intrinsic subunits and cofactors found in mature PSII. Psb27 and Psb28 are well-known assembly factors (Dobáková et al., 2009; Komenda et al., 2012; Mabbitt et al., 2014; Nowaczyk et al., 2012; Roose and Pakrasi, 2008), whereas the additional single transmembrane helix protein (tsl0063), which we named Psb34, has not been described before. The small subunit PsbY, which is known to be loosely bound (Broser et al., 2010), is not resolved in our structure. In addition, PsbJ is not present, as the corresponding gene was inactivated to stall PSII assembly at this specific transition (Nowaczyk et al., 2012).

**Fig. 1:**
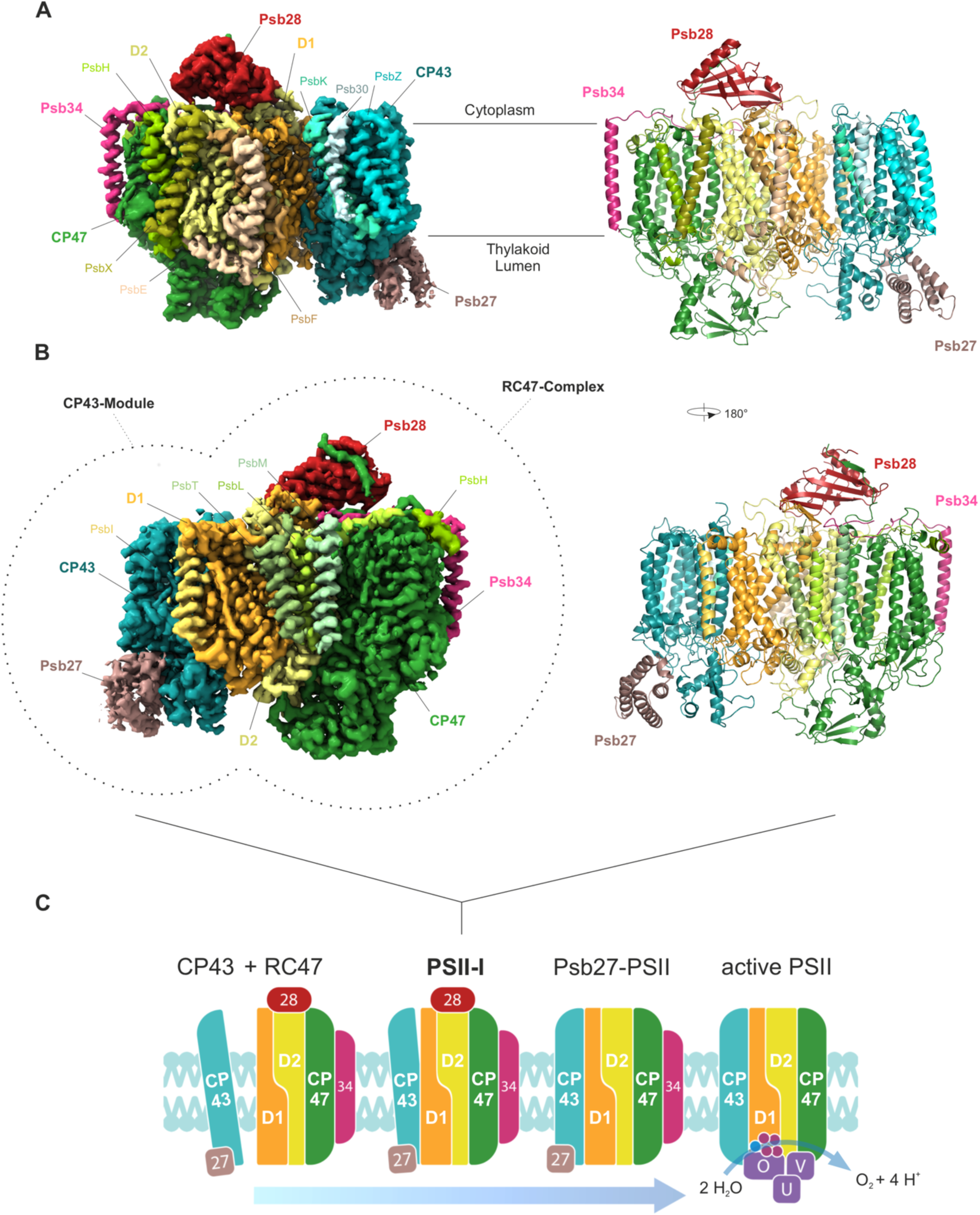
Cryo-EM map of a PSII assembly intermediate (PSII-I) from *T. elongatus*, segmented by subunit. **(A)** 15 PSII subunits and 3 assembly factors are colored and named (PSII subunits: D1, D2, CP43, CP47, PsbE, PsbF, PsbH, PsbI, PsbK, PsbL, PsbM, PsbT, PsbX, PsbZ and Psb30; assembly factors: Psb27, Psb28 and tsl0063, which we named Psb34) (front view). (B) Parts of PSII that originate from the CP43 module (comprised of CP43, Psb27, PsbZ, Psb30 and PsbK) and the RC47 complex are indicated by dashed lines (back view). Schematic model of the PSII assembly process starting with the formation of PSII-I from the CP43 module and RC47. Small PSII subunits were omitted for simplicity.

The two additional maps serve as internal controls. PSII-I’ (2.76 Å) lacks Psb27 but is otherwise comparable to PSII-I; the root mean square deviation (RMSD) of the C_α_ atomic positions between similar subunits of the two complexes is 0.4 Å. Most likely, Psb27 was partly lost during sample preparation. The third cryo-EM map (2.82 Å), which we call PSII-M (for PSII-Monomer), represents a monomeric PSII complex without bound assembly factors. Comparison of our PSII-M structure with a crystal structure of monomeric PSII (Broser et al., 2010) (PDB-ID 3KZI, 3.6 Å) reveals only minimal differences between both structures, with a Cα RMSD of 1.3 Å, which verifies that the structural changes observed in PSII-I are not caused by the deletion of PsbJ.

### Psb34 specifically assists the attachment of the CP43 module to RC47

Our PSII-I structure provides the first identification of the single transmembrane helix protein Psb34 bound to a PSII assembly intermediate (Fig. 2A), which we also confirmed by mass spectrometry (Fig. 2B). Psb34 was probably overlooked previously due to its hydrophobicity and small size. It has a single transmembrane helix that binds to the CP47 antenna protein in close proximity to PsbH (Fig. 2A). Its conserved long N-terminal arm is located at the side and top of the D2 subunit (Fig. 2A). In addition, we independently confirmed the interaction of Psb34 with PSII assembly intermediates by isolation of strep-tagged Psb34 complexes, indicating a specific function of Psb34 in the attachment of CP43 to RC47 (Fig. 2C). Two distinct PSII intermediates were isolated via pulldown of strep-tagged Psb34: the RC47 complex with bound Psb28 and the subsequent PSII intermediate after attachment of CP43 and Psb27 (Fig. 2C). This observation implies that Psb28 is usually released from the PSII intermediate after attachment of CP43, probably after incorporation of PsbJ, as this trigger is missing in the analyzed Δ*psbJ* mutant. Psb34 shows sequence similarity to high-light inducible proteins (HLIPs), which play a role in transient chlorophyll storage and chlorophyll biosynthesis (Komenda and Sobotka, 2016). However, the chlorophyll binding motive is missing in Psb34 (Table S2), suggesting a distinct function for this protein in PSII biogenesis.

**Fig. 2:**
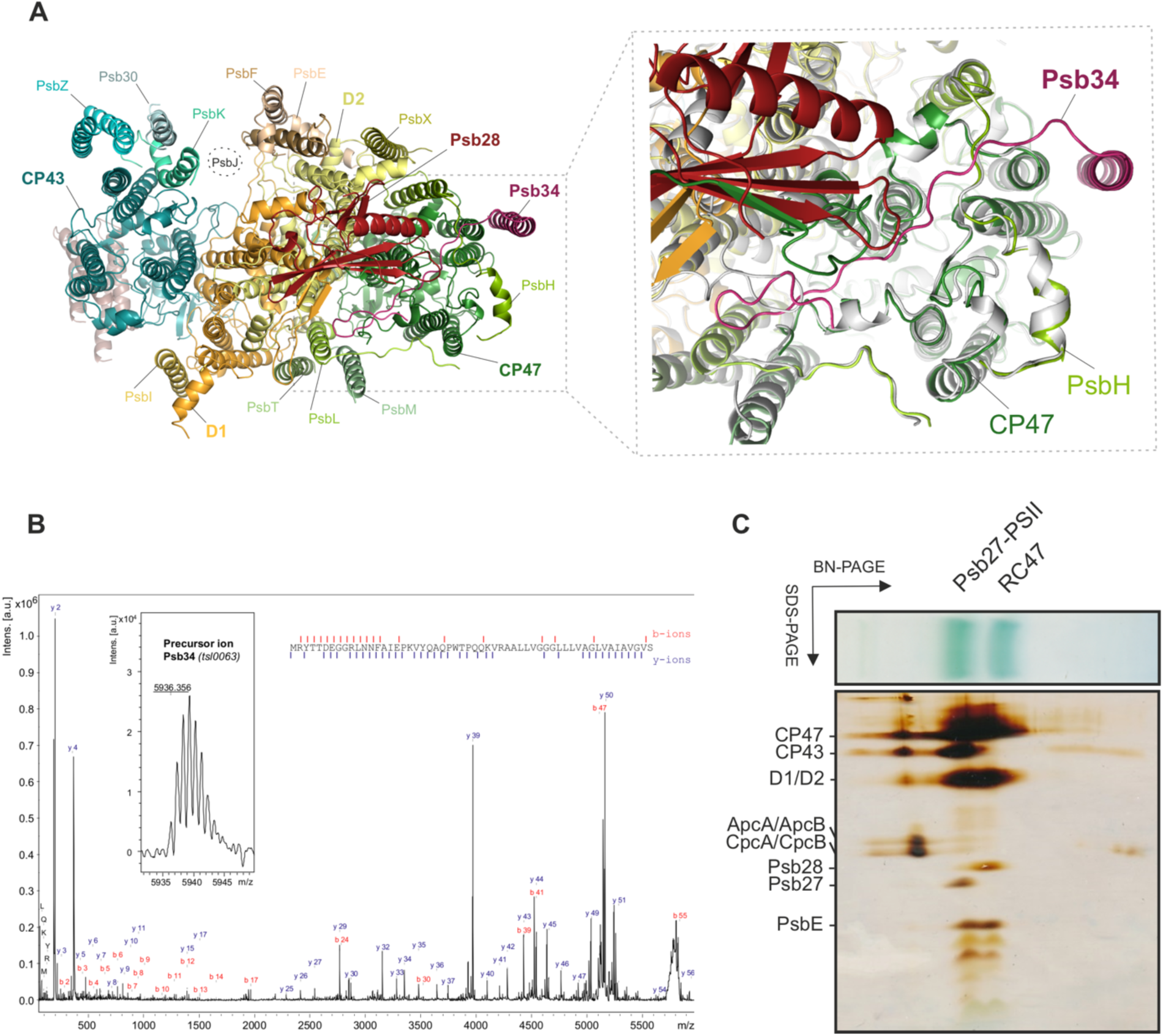
Psb34 binds to RC47 during attachment of the CP43 module. (**A**) Binding site of Psb34 at CP47, close to PsbH (top view), with extended binding of the Psb34 N-terminus along the cytoplasmic PSII surface (dashed box). (**B**) MALDI-ToF analysis of PSII assembly intermediates. Mass spectrum of Psb34 (tsl0063) from the PSII complex (inset) and the fragment spectrum obtained for m/z 5936.356 with annotated b- and y-ion series matching the Psb34 sequence. Observed fragmentation sites are indicated by dashes in the sequence. Mascot score: 171. (**C**) Subunit composition of Psb34-PSII assembly intermediates analyzed by 2D-PAGE.

### Psb28 forms an extended beta hairpin structure that involves the D1 D-E loop and the CP47 C-terminus

Psb28 binds on the cytosolic faces of the D1 and D2 subunits, directly above the Q_B_ binding site (Fig. 3A), which differs from the position that was previously predicted by mass spectrometry (Weisz et al., 2017). Its binding induces the formation of an extended beta-hairpin structure that incorporates the central anti-parallel beta-sheet of Psb28, the C-terminus of CP47 and the D1 D-E loop (Mulo et al., 1997) (Fig. 3A). Binding of Psb28 to the C-terminus of CP47 also imparts a directionality to the assembly process. In the Psb28-free complex (PSII-M), the CP47 C-terminus blocks the Psb28 binding site by interacting with the D1 D-E loop, thus preventing the reverse process and perturbation of active PSII by Psb28. Using nuclear magnetic resonance (NMR) spectroscopy, we performed chemical shift perturbation (CSP) experiments with recombinant Psb28 and a synthetic peptide of the conserved CP47 C-terminus to characterize this interaction in detail and determine the dissociation constant (K_D_) (Fig. 3 and Fig. S4). The CSP measurements indicated significant shifts with a chemical shift difference (Δδ) of more than one standard deviation located at strands β3 and β4 as well at the C-terminal region of Psb28 (Fig. 3C and D). Upon peptide binding, resonances for several residues gradually appeared with increasing peptide concentration, which were line-broadened beyond detection for the free form of Psb28. This observation indicates a less dynamic and more rigid complex structure. This is further supported by the heteronuclear Overhauser effect (NOE) data, which show that the C-terminus of Psb28 becomes rigid from L108 to K112 upon CP47 peptide binding due to creation of an intermolecular β-sheet (Fig. 3E). 2D-lineshape analysis was performed, yielding a K_D_ of 53.92 ± 0.41 µM and a dissociation rate k_off_ of 10.14 ± 0.16 s^-1^, which is consistent with the observed slow-exchange in the NMR spectra (Fig. 3B). The affinity of Psb28 for full-length CP47 and PSII might indeed be even higher due to additional contacts between Psb28 and the D-E loop of D1 (Fig 3A).

**Fig. 3:**
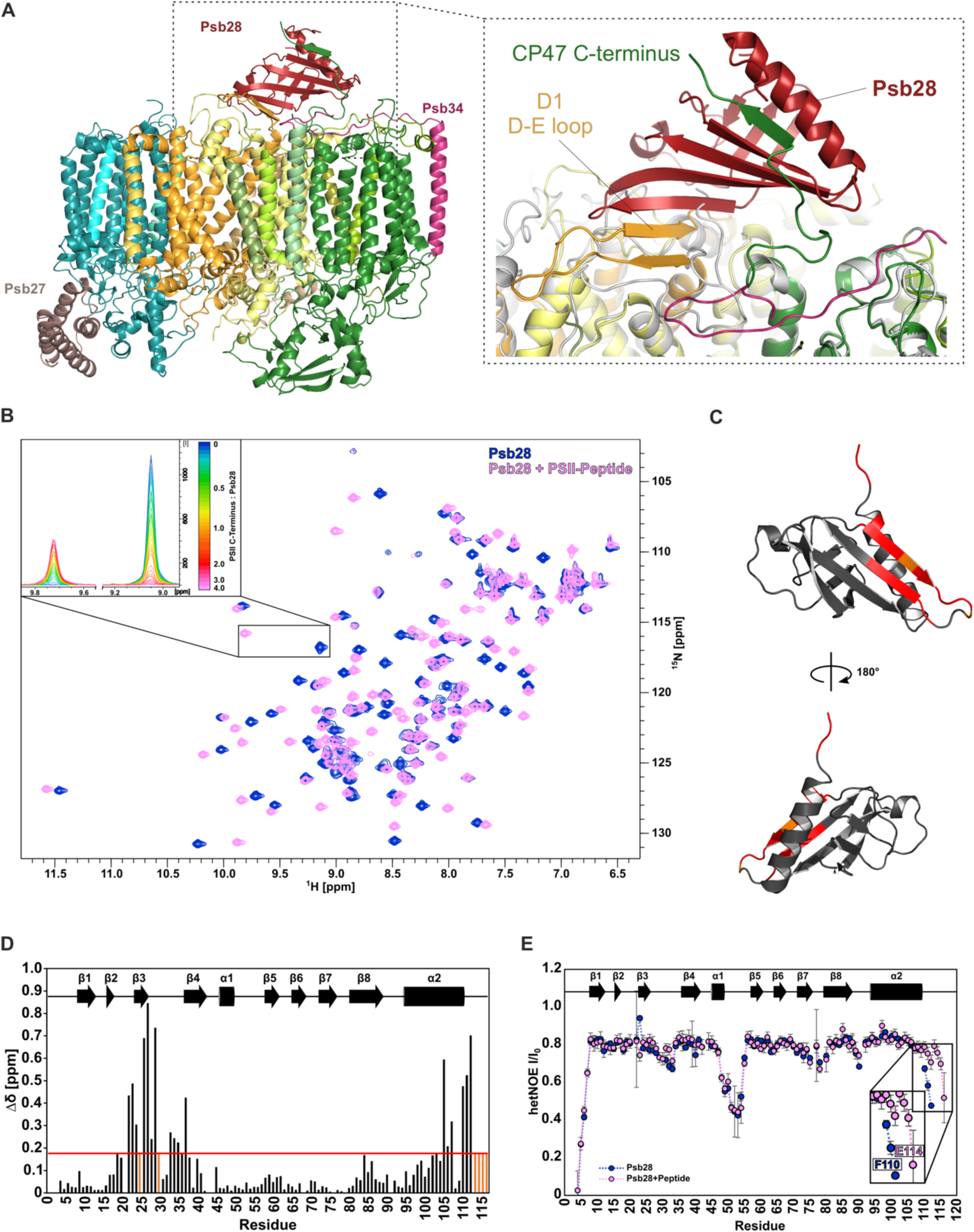
The role of the CP47 C-terminus in binding of Psb28. (**A**). Binding of Psb28 at the cytoplasmic/stromal PSII surface (side view, colors correspond to Fig. 1) and continuation of the central Psb28 beta-sheet by the CP47 C-terminus and the D-E loop of D1 (dashed box). For comparison, mature monomeric PSII (PDB-ID 3KZI) is shown in gray. **(B)** Superimposed 2D ^1^H-^15^N-HSQC spectra of free Psb28 (blue) and Psb28 bound to the C-terminal peptide of CP47 (magenta). Upper left inset: representation of slow exchange behavior for the proton amide resonance of T24, ranging from 126.9 ppm to 128.6 ppm in the ^15^N dimension. **(C)** CSPs of more than one SD projected onto the model representation of Psb28. **(D)** Weighted ^1^H/^15^N chemical shift perturbations observed for Psb28 upon binding to the CP47 peptide. Red line indicates one standard deviation (SD), residues that yield resonances only in the complex form are indicated in orange. **(E)** Backbone ^15^N{^1^H}-heteronuclear NOE of free Psb28 (*blue*) and Psb28 bound to the C-terminal region of the CP47 peptide (*magenta*). Smaller I/I_0_ ratios correspond to regions that exhibit dynamics on the pico- to nanosecond timescale.

### Psb28 binding prevents full association of CP43 and distorts the Q_B_ binding pocket

Binding of Psb28—with support of Psb34—causes major structural perturbations at the PSII acceptor side (Supplementary Movies 1 and 2), which mainly involve the D-E loops of the central D1 and D2 subunits. Comparison of the CP43 structure in PSII-I with that in our PsbJ-free control PSII-M (Fig. 4A-D) or with that in mature monomeric PSII (PDB-ID 3KZI) (Fig. 4C and D) reveals several differences. The CP43 C-terminus is not resolved in PSII-I, probably due to an immature position of the last transmembrane helix of CP43 and an altered conformation of the D1 D-E loop, which may prevent binding of the CP43 C-terminus to the cytoplasmic PSII surface (Fig. 4B). This region is close to the loop between helices D and E of the D2 subunit, which is also altered by binding of Psb28, as clearly shown by movement of D2 Arg233 (Fig. 4B, Fig. S5A and B). After dissociation of Psb28, the CP43 module undergoes a rigid body rotation where it clicks into place (Fig. 4B-D, Supplementary Movie 1), whereas binding of PsbJ and the extrinsic proteins PsbO, PsbV and PsbU during further maturation has very little influence on the CP43 binding position (Fig. 4C and D). The part of PSII that originates from RC47 shows almost no difference between PSII-I and mature PSII (Fig. 4D), except for PsbE, which binds adjacent to PsbJ (Fig. 4C).

**Fig. 4.**
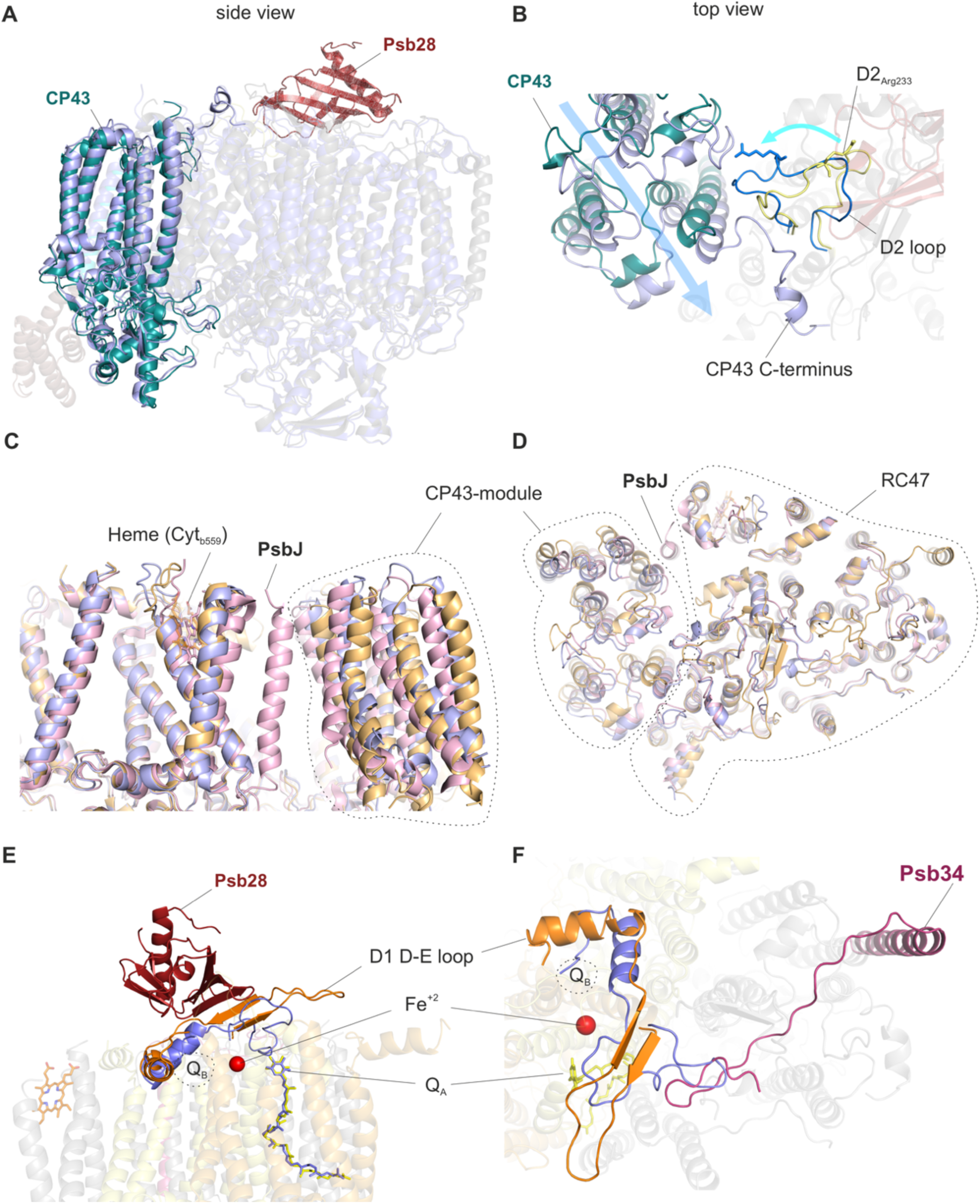
Structural changes of the D1 and D2 D-E loops induced by binding of Psb28 and Psb34. **(A)** Side view of the CP43 antenna protein in PSII-I (*teal*) and the PSII-M control (*light blue*). **(B)** Structural changes between PSII-I and the PSII-M control in the cytoplasmic D2 D-E loop (*yellow*: PSII-I, *blue*: PSII-M) and attachment of CP43 (*teal*: PSII-I, *light blue*: PSII-M control) (top view). Details of the structural changes in the D2 loop are shown in Fig. S5A and B. **(C)** Side view and **(D)** top view of the PSII-I structure (*orange*) compared to the PSII-M control (*light blue*) and mature monomeric PSII (*light red*, PDB-ID 3KZI). **(E)** Side view and **(F)** top view of the Psb28-induced structural changes in the D1 D-E loop (*orange*) and perturbation of the Q_B_ binding site compared to PSII-M (*light blue*), which lacks the assembly factors. Q_A_ is shown in *yellow* (PSII-I) or *light blue* (PSII-M), respectively. See Fig. S5C-H for enlarged views of the Q_A_ and Q_B_ binding site and the adjacent non-heme iron.

Most importantly, the structural changes in the D1 D-E loop may have a direct functional impact on PSII electron transfer (Fig. 4E and F), as this region coordinates several important PSII cofactors. In functional PSII, after charge separation at the reaction center P_680_, electrons are transferred via pheophytin to the bound plastoquinone (Q_A_) and further to mobile plastoquinone (Q_B_). In our PSII-I structure, the Q_A_ site is fully assembled, and a well-resolved Q_A_ molecule is bound (Fig. 4E and F, Fig. S5C and D). The nearby non-heme iron is also already in place in PSII-I (Fig. 4E and F, Fig. S5E and F). The Q_B_ binding site of the PSII-M control is comparable to mature PSII, although it is not occupied by Q_B_ in our preparation (Fig. S5G). In contrast, the Q_B_ binding site of PSII-I is immature due to the Psb28- and Psb34-induced structural changes in the D1 D-E loop (Fig. 4E and F, Fig. S5H). Notably, D1 Phe265, which coordinates the head group of Q_B_ in mature PSII, is clearly at a different position (Umena et al., 2011) (Supplementary Movie 2).

### Binding of Psb28 protects PSII during biogenesis

A more detailed analysis of the structural environment close to the Q_A_/Q_B_ binding sites revealed differences in the coordination and the hydrogen-bond network of the adjacent non-heme iron, which also indicate functional consequences for PSII electron transfer and charge recombination processes. In mature PSII, the non-heme iron is coordinated by four histidine residues and bicarbonate as the fifth ligand (Fig. 5A and C), whereas in PSII-I, the bicarbonate molecule is replaced by the E241 side-chain of D2 (Fig. 5B and D, Fig. S5E and F, Supplementary Movie 3). Other residues, including D1 E244 and Y246, which bind to the bicarbonate molecule in mature PSII (Fig. 5A), are also displaced in PSII-I due to the conformational change of the D1 D-E loop (Fig. 5B, Fig. S5E and F, Supplementary Movie 3). Binding of bicarbonate is important for PSII efficiency (Eaton-Rye and Govindjee, 1988), as it lowers the redox potential of (Q_A_/Q_A_^-^) to favor forward electron transport (Allen and Nield, 2017; Brinkert et al., 2016). If charge recombination occurs, the lower redox potential favors indirect charge recombination via P^•+^/Pheo^•-^. This back reaction yields triplet chlorophyll and subsequently singlet oxygen (Brinkert et al., 2016), a highly oxidizing species. Changes in the redox potential of (Q_A_/Q_A_^-^) have been proposed to tune the efficiency of PSII depending on the availability of CO_2_ as the final electron acceptor and thereby protect PSII under low CO_2_ conditions (Brinkert et al., 2016). Therefore, we used flash-induced variable fluorescence to measure electron transfer in the PSII-I assembly intermediate and inactivated PSII, both of which lack a functional OEC (Fig. 5E, Fig. S6A and B). The fast component is assigned to PSII centers with fast reoxidation of Q_A_^-^ by properly bound Q_B_, the middle component is caused by PSII complexes with inaccurately bound Q_B_, and the slow component is associated with centers that do not contain Q_B_ and instead reoxidize Q_A_^-^ through charge recombination with the Mn_4_CaO_5_ cluster (Vass et al., 1999). Fully functional PSII showed typical Q_A_^-^ reoxidation, which is primarily due to fast electron transfer to Q_B_ (Fig. 5E, blue trace). Addition of DCMU blocks electron transfer to Q_B_ in active PSII, thereby promoting slow S_2_Q_A_^-^ charge recombination (Fig. S6B, blue trace). Removal of the OEC increases the Q_A_ redox potential (Johnson et al., 1995) and promotes very slow Q_A_^-^Tyr_D_^+^ recombination (Fig. 5E, black trace) (Johnson et al., 1994), which is influenced only minorly by binding of DCMU (Fig. S6B, black trace). PSII-I shows a different behavior (Fig. 5E, Fig. S6B red trace); ∼60% of the PSII-I centers decay within 1 s, whereas ∼40% decay in PSII (-OEC). To determine whether the replacement of bicarbonate by glutamate affects the energetics of the redox couple Q_A_/Q_A_^-^, we measured the formation of ^1^O_2_ by EPR spectroscopy using the spin probe TEMPD. The data clearly show that ^1^O_2_ formation is reduced by ∼30% in PSII-I compared to inactivated PSII (-OEC), which contains bicarbonate (Fig. 5F).

**Fig. 5:**
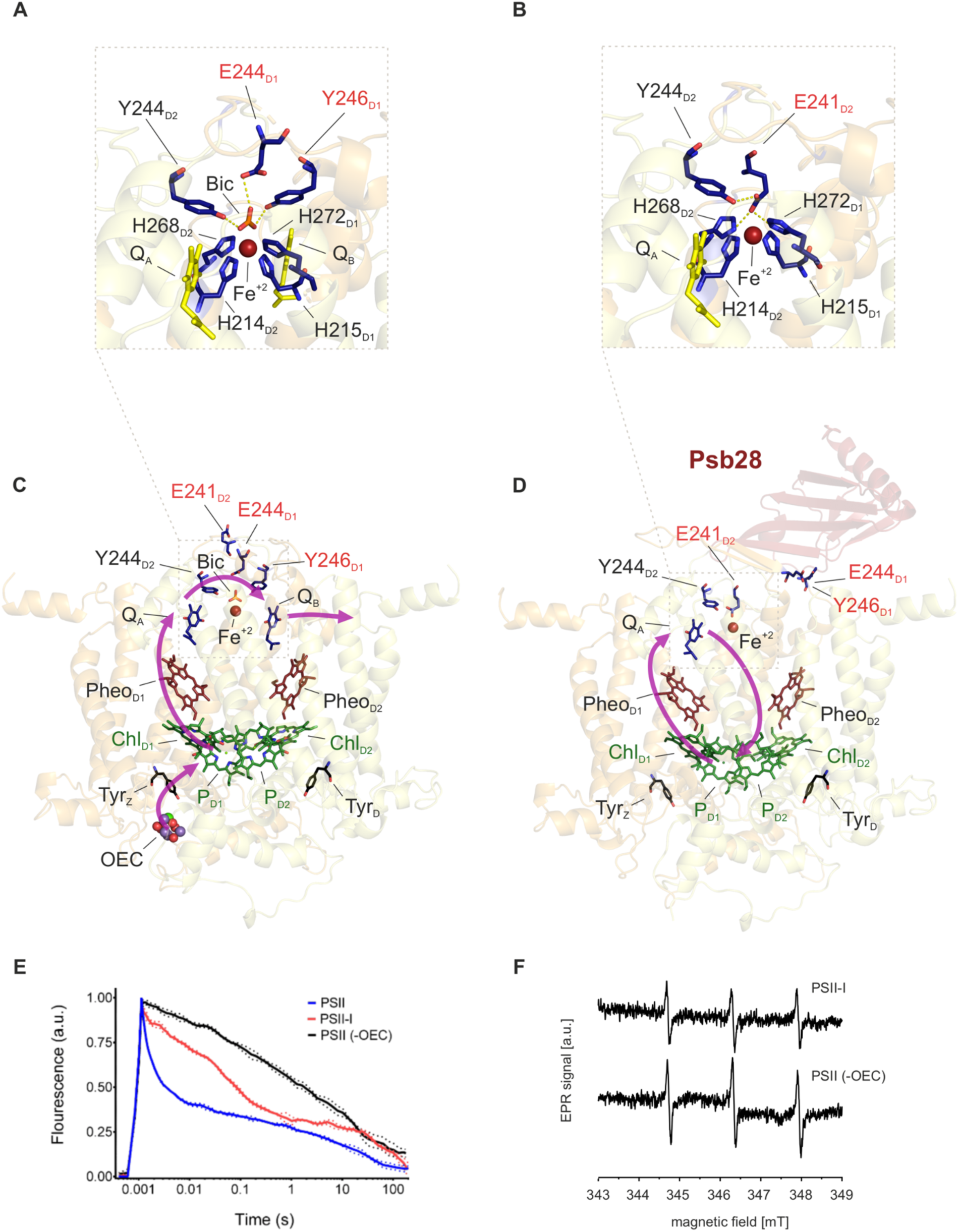
Binding of Psb28 displaces bicarbonate as a ligand of the non-heme iron and protects PSII from damage. (**A)** The electron transfer from PQ_A_ to PQ_B_ is coordinated by the non-heme iron (Fe^2+^), with the binding of bicarbonate (Bic) serving as a regulatory mechanism (Brinkert et al., 2016) in mature PSII (PDB-ID 3WU2). **(B)** Binding of Psb28 to the PSII-I assembly intermediate induces a conformational change in the cytoplasmic D2 D-E loop, where the side chain of Glu241 replaces bicarbonate as a ligand of the non-heme iron. The respective fits of the non-heme iron binding sites are shown in Fig. S5E and F. A similar coordination is also found in non-oxygenic bacterial reaction centers (Stowell et al., 1997) (Fig. S6C). **(C)** Electron transfer (purple arrows) in mature PSII. Light-induced charge separation at the reaction center chlorophylls (P_D1_, P_D2_, Chl_D1_, Chl_D2_) leads to electron transfer via pheophytin (Pheo_D1_) and plastoquinone A (Q_A_) towards Q_B_. The electron gap at the reaction center is filled by the oxygen evolving complex (OEC). (**D**) Reoxidation of Q_A_^-^ by direct and safe charge recombination is favored in the PSII assembly intermediate, as indicated by the purple arrows. (E) Flash-induced fluorescence decay of PSII. Blue lines represent active PSII, black and red represent PSII without functional OEC and PSII-I respectively. Dotted corridors depict SD (*n* = 3). **(F)** The protective role of Psb28 binding was further confirmed by EPR spectroscopy using the spin probe TEMPD, which is specific for ^1^O_2_, the major reactive oxygen species in PSII generated by triplet chlorophyll (^3^P). Inactivated PSII without functional Mn_4_CaO_5_ cluster (-OEC) was used as control.

### Psb27 binds in a remote position to loop E of CP43 at the luminal PSII surface

Psb27 binds to the luminal side of the PSII complex, adjacent to loop E of the CP43 subunit (Fig. 6A). In contrast to previously proposed models (Cormann et al., 2016; Liu et al., 2011a), the binding site of Psb27 has little overlap with the binding sites of the extrinsic subunits (PsbO, PsbV and PsbU) and has at least no direct impact on the Mn_4_CaO_5_ cluster binding site (Fig. 6A and B). Instead, Psb27 is bound at a remote position that might be occupied by CyanoQ in the mature complex (Wei et al., 2016). This localization of Psb27 does not support previous functional models in which bound Psb27 prevents the binding of the extrinsic subunits or plays a direct role in Mn_4_CaO_5_ cluster assembly (Cormann et al., 2016; Nowaczyk et al., 2006). However, Psb27 might stabilize loop E of CP43 in the unassembled state and facilitate its binding to the D1 subunit. This is of particular importance, as loop E of CP43 provides Arg345 and Glu342, two ligands of the Mn_4_CaO_5_ cluster in mature PSII (Fig. 6B, dashed box). Moreover, in the Psb27-bound state (PSII-I), the D1 C-terminus, which is directly involved in coordination of the Mn_4_CaO_5_ cluster (Umena et al., 2011), is bound away from the cluster (Fig. 6C, Fig. S7). Thus, our PSII-I structure reveals not only how the Psb27 protein binds to CP43 and thus stabilizes it prior to attachment, but also indicates an indirect role for CP43 in maturation of the oxygen evolving cluster that is consistent with functional data from previous studies (Grasse et al., 2011; Komenda et al., 2012; Roose and Pakrasi, 2008).

**Fig. 6:**
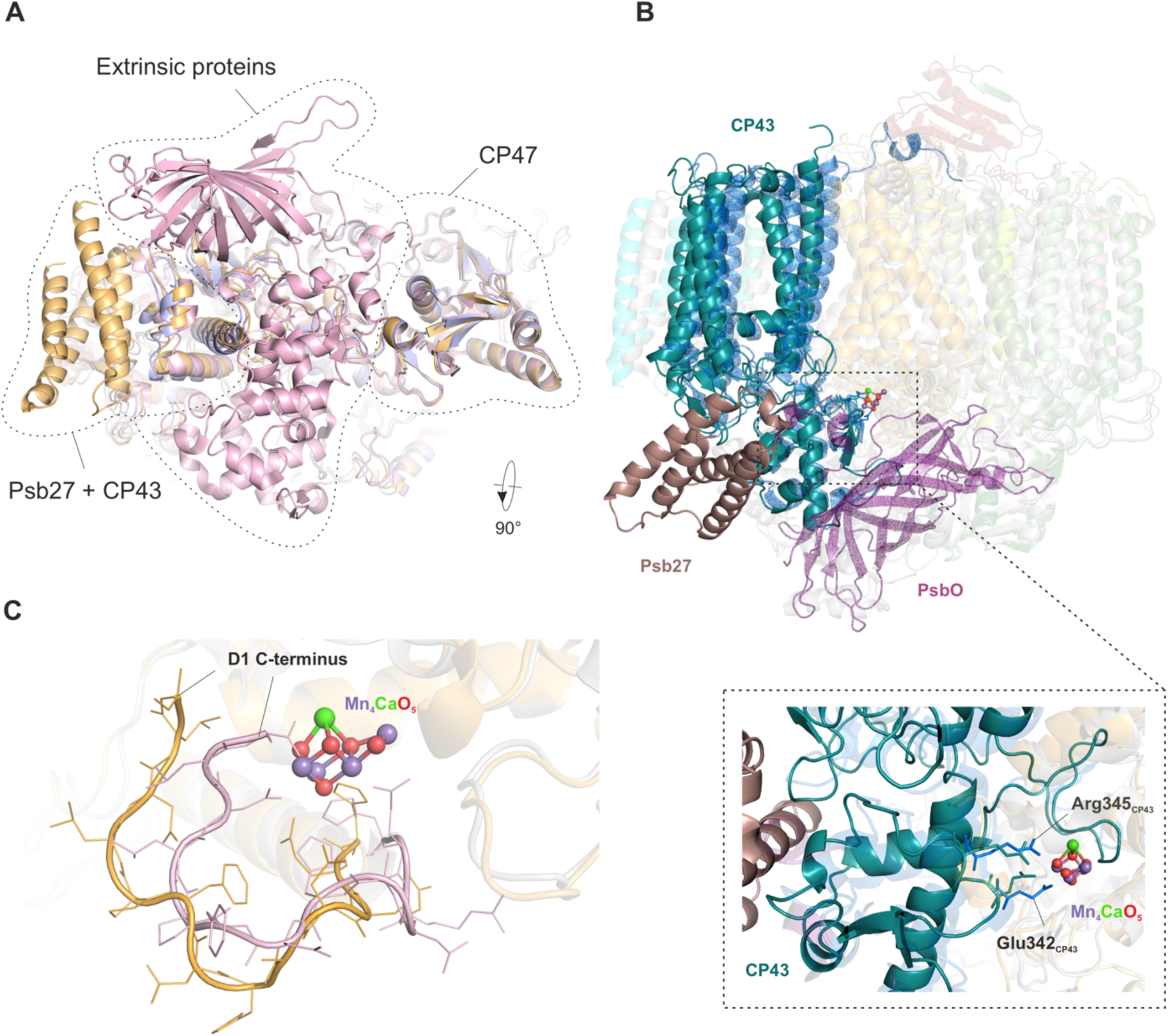
The role of Psb27 in Mn_4_CaO_5_ cluster assembly. (**A**) Bottom view of the luminal PSII surface for PSII-I (*orange*), the PSII-M control (*light blue*) and mature monomeric PSII (PDB-ID 3KZI) *(light red*). **(B)** Side view of CP43 (*teal*) and Psb27 (*brown*) in PSII-I, as well as of CP43 (*blue)* and PsbO (*purple*) in mature monomeric PSII (PDB-ID 3KZI). Dashed box: CP43 E loop with residues Arg345 and Glu342 (shown as sticks), which are involved in coordination of the Mn_4_CaO_5_ cluster. We changed the numbering of CP43 residues due to a corrected N-terminal sequence (www.UniProt.org). The residues correspond to Arg357 and Glu354 in previous publications. The high-resolution structure of the Mn_4_CaO_5_ cluster is taken from Umena et al. 2011 (PDB-ID 3WU2). **(C)** Position of the D1 C-terminus in PSII-I (*orange*) and mature monomeric PSII (PDB-ID 3KZI) (*light red*).

### The immature Mn_4_CaO_5_ cluster binding site of PSII-I contains a single, positively charged ion

The unique Mn_4_CaO_5_ cluster is a key feature of PSII that splits water into oxygen and protons. However, our PSII-I complex does not show any oxygen-evolving activity, suggesting that the oxygen evolving complex (OEC) is not fully assembled. In mature PSII, the Mn_4_CaO_5_ cluster is submerged in the complex and additionally capped by the extrinsic subunits PsbO, PsbU and PsbV (Fig. 6A and B). In our PSII-I structure, these subunits are absent, which leaves two parts of the CP43 E-loop (residues 320-327 and 397-404) in a flexible conformation, exposing the binding site of Mn_4_CaO_5_ cluster to the lumen. There is no strong density feature at this position that would correspond to the fully assembled metal-redox cofactor. Thus, our PSII-I structure provides a model for an immature OEC. By comparing our structure with the high-resolution crystal structure of mature PSII (Umena et al., 2011)(PDB-ID 3WU2) provides insights into the first-steps of OEC biogenesis (Fig. 7).

**Figure 7:**
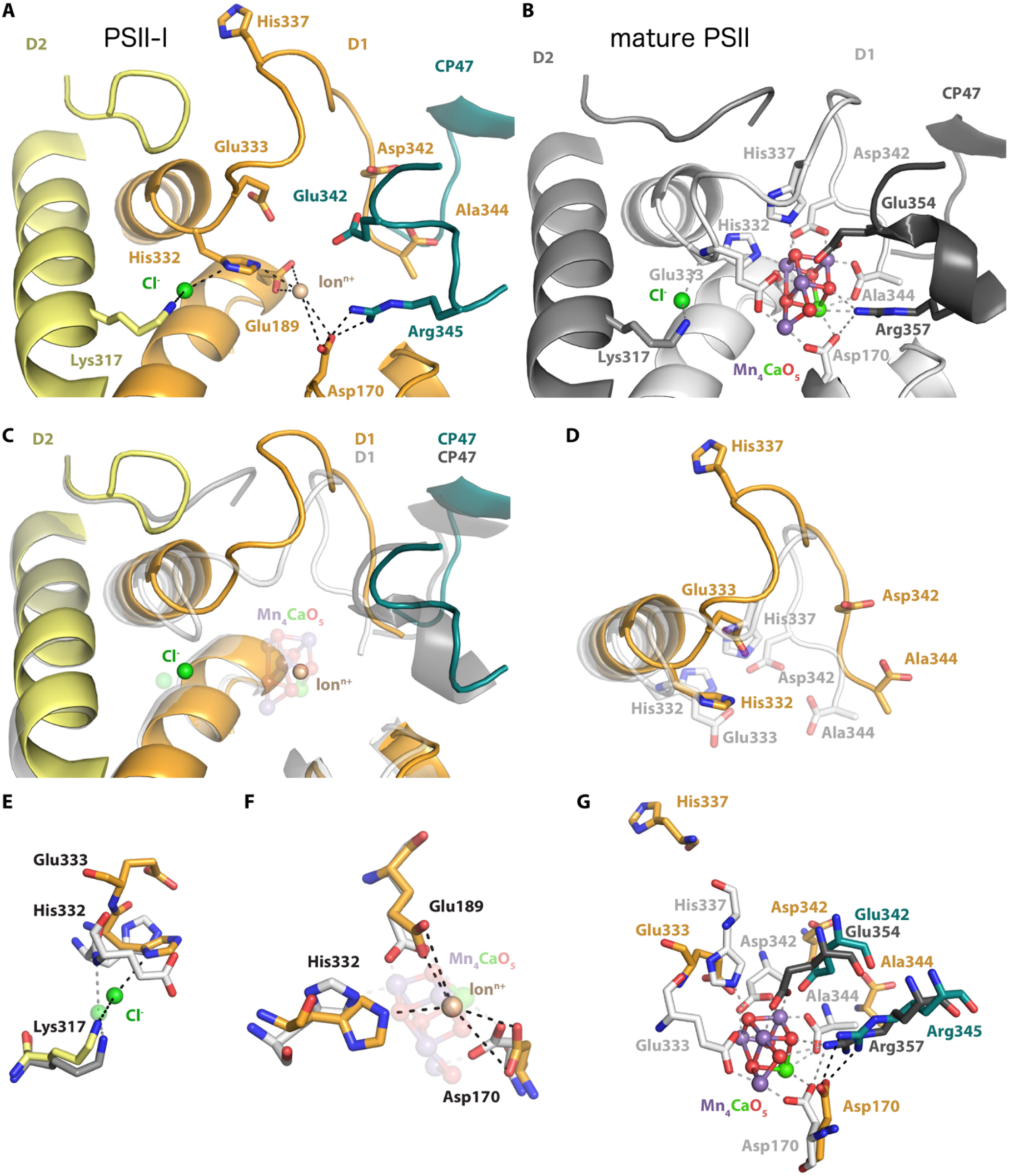
Conformational changes within the active site of the Mn_4_CaO_5_ cluster. The Mn_4_CaO_5_ cluster performs PSII’s unique water-splitting reaction. (**A**) The active site of the Mn_4_CaO_5_ cluster is resolved within our PSII-I structural model but is not yet oxygen-evolving. (**B**) Crystal structure of the oxygen-evolving, mature PSII (PDB-ID 3WU2, resolution 1.9 Å). (**C**) Overlay of both structures, illustrating significant differences in the backbone conformation of the D1 and D2 C-terminal tails. (**D**) Accompanying side chain rearrangements of the D1 C-terminus. The Cl^-^ (**E**), Ion^+^ (**F**) and Mn_4_CaO_5_ (**G**) cluster coordination partners are compared in detailed. The validation of the fit to density for the structural details shown here is provided in Figure S7.

The D1 C-terminus is one of the key features for the formation of the of OEC, as it provides several essential charged residues that are responsible for coordination of the chloride ion and the Mn_4_CaO_5_ cluster (Fig. 7A, B and D). The density for these C-terminal residues is weak in our PSII-I map, but traceable (Fig. S7A), indicating a flexibility that confirms the absence of the OEC. Compared to the mature complex, the last 12 residues of the C-terminal tail of D1 would need to undergo significant conformational changes to bring the side chains of Glu333, His337, Asp342, and the Ala344 C-terminus into the correct position to coordinate the Mn_4_CaO_5_ cluster (Supplementary Movie 4).

Moreover, we identify a clearly visible density at the position of the chloride ion, which is coordinated by Lys317 (D2) and the hydrogen atom of the backbone nitrogen of Glu333 (D1) in mature PSII (Fig. 7B and E). Despite the similar position, the Cl^-^ is coordinated by the nitrogen atom of the ring of adjacent His332 (D1) in PSII-I (Fig. 7A and E, Fig. S7D). Surprisingly, we identified another density in the area where the Mn_4_CaO_5_ cluster is located in mature PSII (Fig. 7A-C and F, Fig. S7C). However, this density is not large enough to reflect the whole cluster. Based on its size and interaction partners (Fig. 7F), it corresponds to one positively charged ion. In the structural context, this ion is most likely Mn^2+^, but it could also be Ca^2+^ or any other positively charged ion.

This ion is coordinated by the side chains of D1 Asp170, Glu189, and His332, which are already in similar positions compared to mature PSII. Glu342 and Arg345 of CP43, which are both involved in the Mn_4_CaO_5_ cluster coordination, are also already pre-positioned through the interaction between Arg345 with D1 Asp170 (Fig. 7G). However, there are still significant conformational changes necessary for the transition from PSII-I to mature PSII, as highlighted in Figure 7D and G, as well as in Supplementary Movie 4. The D1 C-terminal tail must bring the side chains of Glu333, His337 and Asp342, as well as the C-terminus of Ala344, into correct alignment to coordinate the Mn_4_CaO_5_ cluster. In addition, the C-terminal tail of D2 needs to flip towards the D1 C-terminus (Fig. 7C, Fig. S7B, Supplementary Movie 4). In summary, PSII-I is characterized by only one positive charged ion bound instead of the complete Mn_4_CaO_5_ cluster, resulting in significantly different conformations of the D1 and D2 C-termini compared to the structural model containing a mature Mn_4_CaO_5_ cluster. However, the PSII-I structure seems to be prepared to accept the Mn_4_CaO_5_ cluster, as indicated by the above described similarities in side chain positioning.

## Discussion

PSII biogenesis is a complex process that requires the action of specific assembly factors. These auxiliary proteins are not present in the mature complex and interact only transiently with specific subunits or preassembled PSII intermediates. Although more than 20 factors have been identified and allocated to specific transitions, their precise molecular function in PSII assembly remains elusive in almost all cases. Our study provides the first detailed molecular insights into the function of PSII assembly factors Psb27, Psb28 and Psb34, which are involved in an important transition prior to activation of the OEC. The determined binding positions of Psb27 and Psb28, which are two of the most studied PSII assembly factors, disprove all previous Psb27 and Psb28 binding models and exclude Psb27 from direct involvement in OEC maturation (Cormann et al., 2009; Cormann et al., 2016; Fagerlund and Eaton-Rye, 2011; Liu et al., 2013; Liu et al., 2011a; Michoux et al., 2012; Weisz et al., 2016).

Binding of Psb28 and Psb34 to the cytoplasmic side of PSII induces large conformational changes in the D1 D-E loop (Fig. 4), which has been identified previously as an important location for PSII photoinhibition and D1 degradation (Kettunen et al., 1996; Mulo et al., 1998). Structural changes observed in the PSII-I Q_B_ binding pocket and coordination of the non-heme iron suggest a functional impact on PSII electron transfer to protect the immature complex until water splitting is activated. In particular, D2 Glu241 replacing bicarbonate as ligand of the non-heme iron by suggests a regulatory role, as binding of bicarbonate was proposed to tune PSII efficiency by changing the redox potential of (Q_A_/Q_A_^-^) (Allen and Nield, 2017; Brinkert et al., 2016). As a functional consequence, PSII-I generates less singlet oxygen compared to inactive PSII (Fig. 5F).

Interestingly, the coordination of the non-heme iron in PSII-I resembles that in non-oxygenic bacterial reaction centers (BRCs) (Stowell et al., 1997) (Fig. S6C). In BRCs, the fifth ligand of the non-heme iron is provided by E234 of the M subunit (Wang et al., 1992), and mutagenesis of this residue induces changes in the free energy gap between the P^•+^/Q_A_^•-^ radical pair (Cheap et al., 2009). These findings indicate that the environment of the non-heme iron is important for regulation of forward electron transfer to Q_B_ versus charge recombination (Allen and Nield, 2017). Therefore, we speculate that during biogenesis, PSII switches to a mechanism that usually operates in non-oxygenic bacterial reaction centers.

The Psb27-bound and -unbound structures do not differ substantially (Fig. 6B), suggesting a rather subtle action in PSII biogenesis. Previous work demonstrated that Psb27 is already bound to free CP43 (Komenda et al., 2012), where it might protect free CP43 from degradation or stabilize the E-loop in a specific conformation to chaperone the subsequent association with the RC47 complex. This step is crucial for preparing the binding site of the Mn_4_CaO_5_ cluster, as the CP43 E-loop provides two ligands of the cluster. Further OEC assembly is a multistep process that requires a functional upstream redox chain for the oxidation of Mn^2+^ to build up the cluster’s µ-oxo bridges between the manganese atoms (Bao and Burnap, 2016; Becker et al., 2011; Cheniae and Martin, 1971; Dasgupta et al., 2008). The mechanistic and structural details of this photoactivation process are not yet understood. In the consensus ‘two quantum model’, a single Mn^2+^ ion bound to the high-affinity site (HAS) is oxidized to Mn^3+^. This initiating light-dependent step is followed by a slow light-independent phase and further fast light-dependent steps in which the remaining Mn^2+^ ions are oxidized and incorporated. Understanding the light-independent slow phase is key to unraveling the mechanism of photoactivation.

Previous structural studies aimed to obtain mechanistic insights into the dark-rearrangement by removing the Mn_4_CaO_5_ cluster from fully assembled PSII, either by depleting it directly from PSII crystals by chemical treatment (Zhang et al., 2017) or by cryo-EM single particle analysis in manganese- and calcium-free buffer (Gisriel et al., 2020). The X-ray structure was indeed missing the Mn_4_CaO_5_ cluster, but the D1 C-terminus followed mostly the same trajectory as found in the mature PSII-dimer structure. The authors suggested that the D1 C-terminus might not rearrange during Mn_4_CaO_5_ cluster assembly. However, the crystal structure was dimeric and still had the extrinsic subunits PsbO, PsbU, and PsbV bound. It is known that these subunits are typically not associated with Mn_4_CaO_5_ cluster-depleted PSII. Thus, the structure might be artificially stabilized by crystal packing forces. The cryo-EM structure, on the other hand, revealed a monomeric PSII that lacks extrinsic subunits and the Mn_4_CaO_5_ cluster (Gisriel et al., 2020). This structure is more similar to our PSII biogenesis intermediate PSII-I, as PsbY, PsbZ and PsbJ are also missing. The PsbJ subunit is surprising; it is an integral subunit of PSII and should not be easily detached, yet it is missing from this structure and we deleted it to stabilize our PSII-I complex. These observations might indicate a more specific and regulatory role of PsbJ in PSII biogenesis. Additionally, the D1 C-terminus is disordered in this previous cryo-EM structure, and the authors suggest that the dark-rearrangement involves a transition from a disordered to an ordered state.

Our structure now reveals the fate of the D1 C-terminus with the assembly factor Psb27 bound. The D1 C-terminus follows a different trajectory compared to the mature PSII. Thus, we provide structural evidence that the slow dark-rearrangement involves a conformational change of the D1 C-terminus rather than the previously proposed disorder-to-order transition after initial photoactivation (Gisriel et al., 2020). Compared to mature PSII, twelve residues of the D1 C-terminal tail must undergo significant conformational changes to bridge the side chains of Glu333, His337 and Asp342, as well as to bring the C-terminus of Ala344 in the correct position to coordinate the Mn_4_CaO_5_ cluster (Fig. 6C and 7D, Supplementary Movie 4), which is consistent with previous models (Dasgupta et al., 2008; Kolling et al., 2012; Zaltsman et al., 1997). We also identified a single positively charged ion in our PSII-I structure, coordinated by Asp170, Glu189 and His332 of D1 (Fig. 7F), at the position of the Mn_4_CaO_5_ cluster of mature PSII. This binding site most likely corresponds to the long-sought single high-affinity site (HAS), where the first Mn^2+^ binds prior to the first photoactivation step in OEC biogenesis (Nixon and Diner, 1992). However, we cannot exclude binding of Ca^2+^, which was shown to bind with a much lower affinity (Kolling et al., 2012; Tyryshkin et al., 2006), or any other positive charged ion at this position. Nevertheless, Asp170 has been identified as the most critical residue for the HAS (Campbell et al., 2000; Cohen et al., 2007), which supports our hypothesis. Further photoactivation steps occur presumably after cooperative binding of calcium and manganese. The binding of the extrinsic subunit PsbO, potentially after release of Psb27 and maturation of the WOC, is the next step of the PSII assembly line *in vivo*, which leads to the next unsolved question in PSII biogenesis: what triggers the release of an assembly factor? For Psb27, its detachment might be promoted by the binding of PsbO, as their binding sites partially overlap.

Membrane protein complexes play a fundamental role in bioenergetics to sustain and proliferate life on Earth. They drive the light-to-chemical energy conversion in photosynthetic organisms and are essential for energy supply in heterotrophs. These highly complex molecular machines are assembled from numerous single proteins in a spatiotemporally synchronized process that is facilitated by a network of assembly factors. These auxiliary proteins are the key players of Nature’s assembly lines. Our PSII-I cryo-EM structure reveals the first molecular snapshot of PSII biogenesis and, accompanied by our spectroscopic and biochemical analyses, provides clear mechanistic insights into how three assembly factors (Psb27, Psb28 and Psb34) coordinate the stepwise construction of this powerful catalyst of life.

## Supplementary Figures

**Figure S1:**
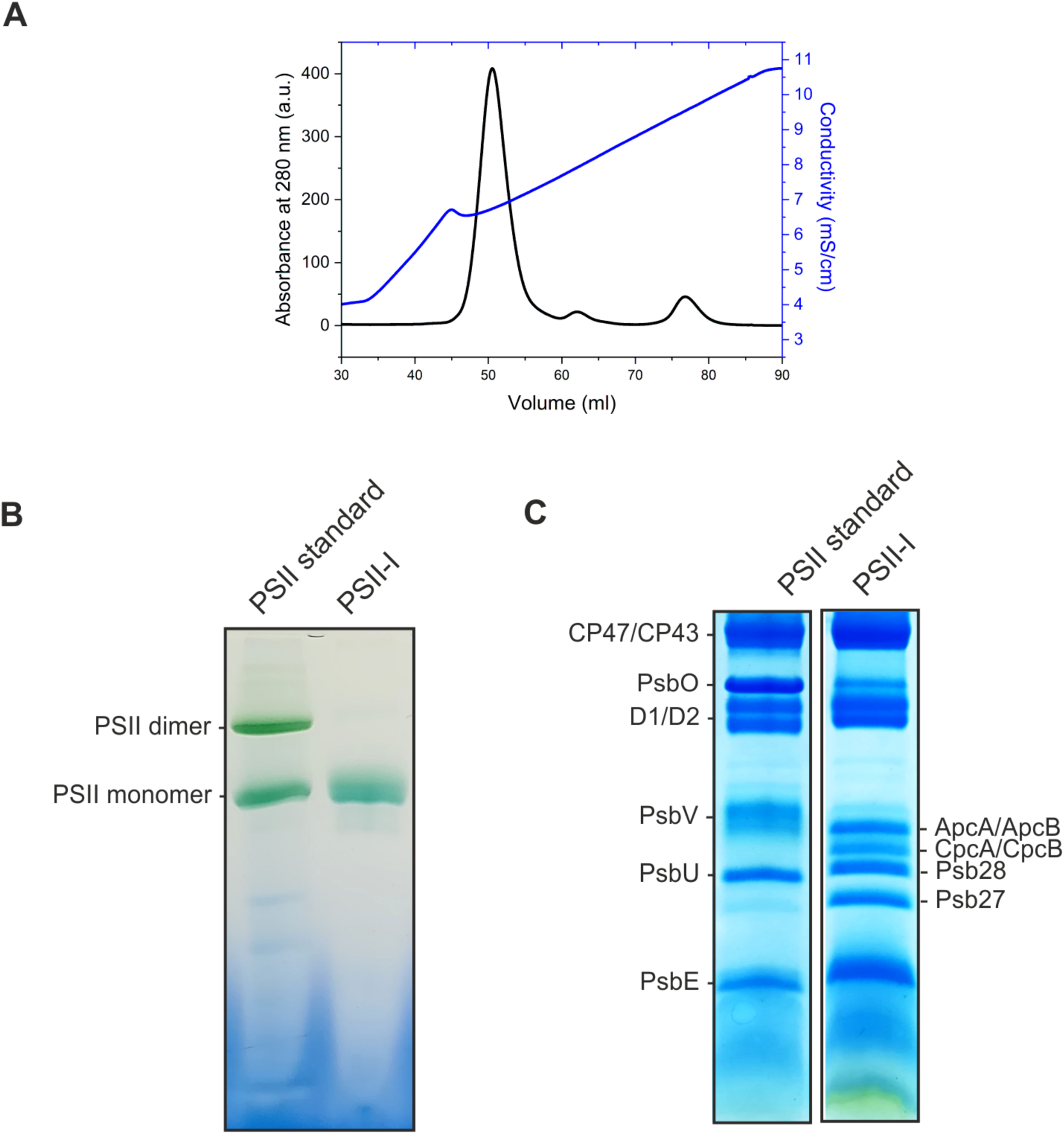
Purification of *T. elongatus* PSII assembly intermediates. **(A)** Streptactin-affinity purification of PsbC-TS PSII assembly intermediates. A_280_ (black) and percentage of elution buffer (blue) plotted against volume. **(B)** Oligomeric state of PSII-I analyzed by Blue-native PAGE. **(C)** Subunit composition of PSII-I analyzed by SDS-PAGE.

**Figure S2:**
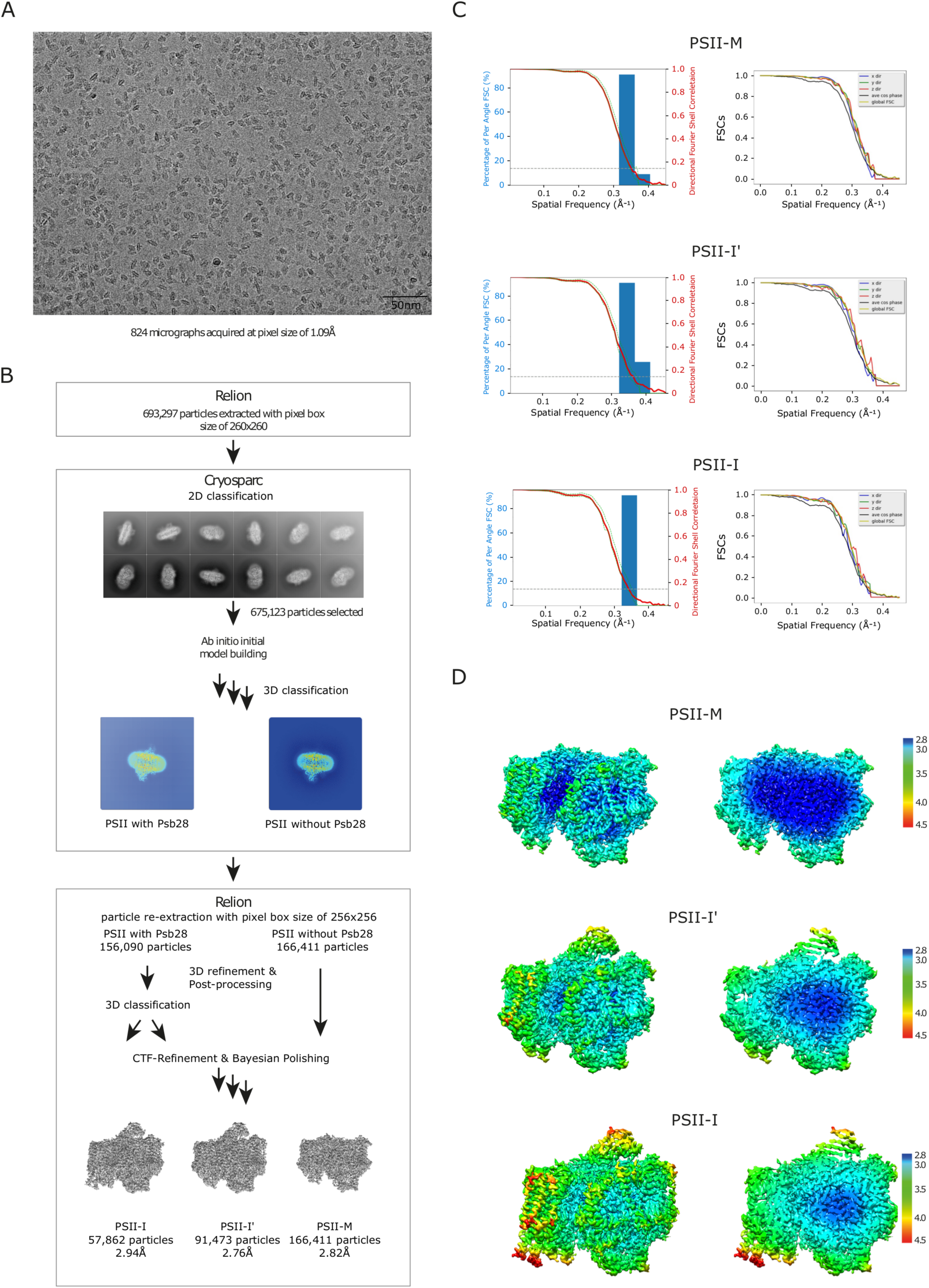
Cryo-EM data collection and processing scheme. **(A)** Example micrograph acquired on a K3 detector (Gatan, 5760×4092 pixel) showing particle distribution on Quantifoil 2/1 grid-holes. **(B)** Schematic of the image processing workflow. After initial particle extraction with RELION, selected particles were used for 2D classification. Selected classes were used for ab initio model building and 3D-classification in Cryosparc. Selected 3D classes were refined and post-processed in RELION. The 3D class containing density for Psb28 was further classified using a mask around the suspected area of Psb27, resulting in densities with and without Psb27. Finally, particles of the resulting three densities were subjected to Bayesian polishing and models were refined. **(C)** Plots showing the global (left plot) and directional (right plot) resolution for each of the three densities obtained in B, calculated using the “Remote 3DFSC Processing Server” web interface (Tan et al., 2017). For PSII-M, PSII-I’ and PSII-I a sphericity score of 0.979, 0.983 and 0.980 and c global resolution (FSC-threshold = 0.143) of 2.82A, 2.76A and 2.94A was calculated, respectively. **(D)** Local resolution as calculated by RELION mapped on the refined densities (left: surface view, right: cut-open view of central section).

**Figure S3:**
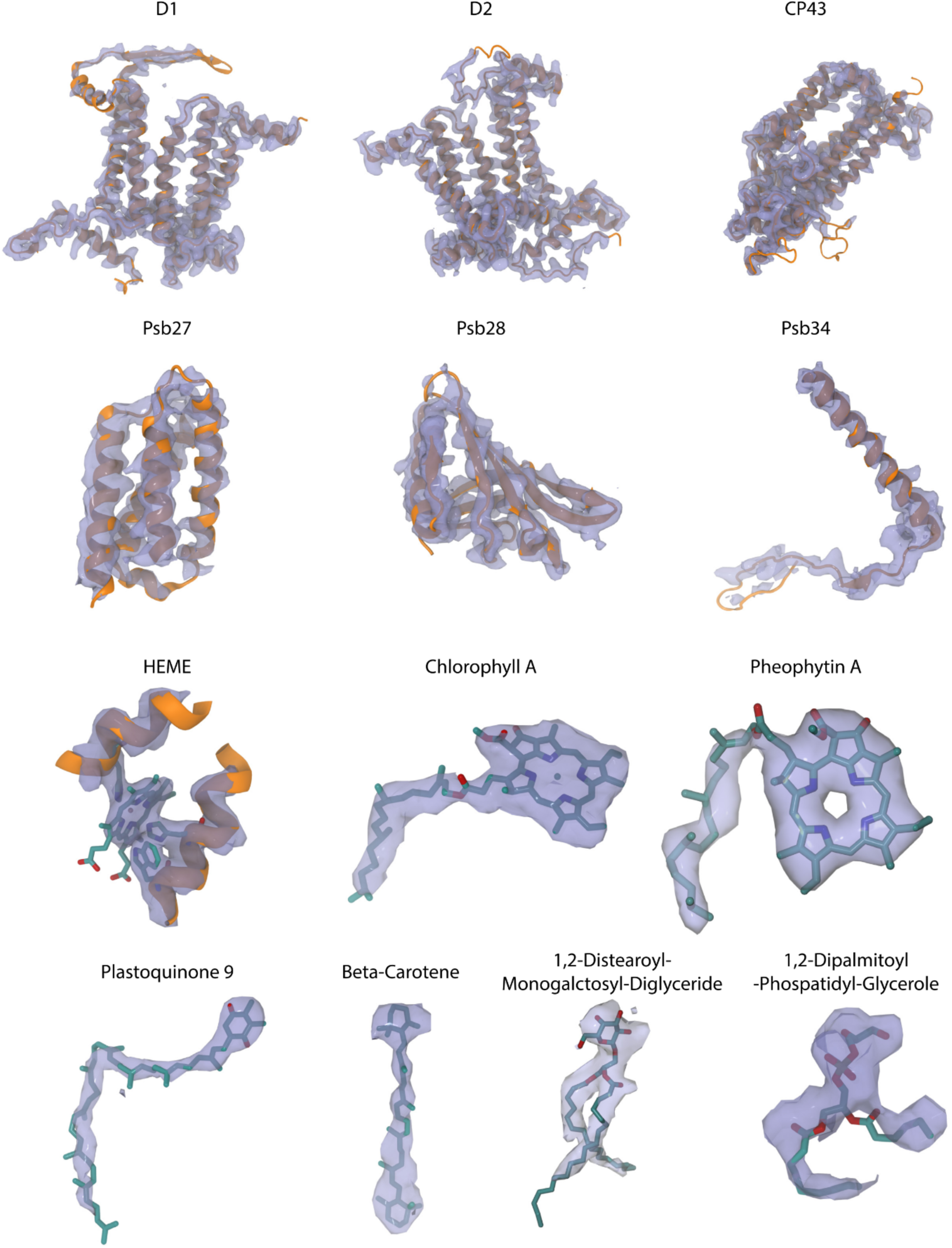
Comparison between the structural model and density for PSII-I. Fit of the structural model into the map density (transparent blue) for selected subunits and cofactors of PSII-I. The resolution is high enough for a reliable assignment of the cofactors and confident modelling of the subunits. The overall fit of the model to the density of PSII-I’ and PSII-M is comparable to the PSII-I examples shown here. The model quality of fit to density of the functionally decisive structural regions and side chains is shown in Figure S5.

**Figure S4:**
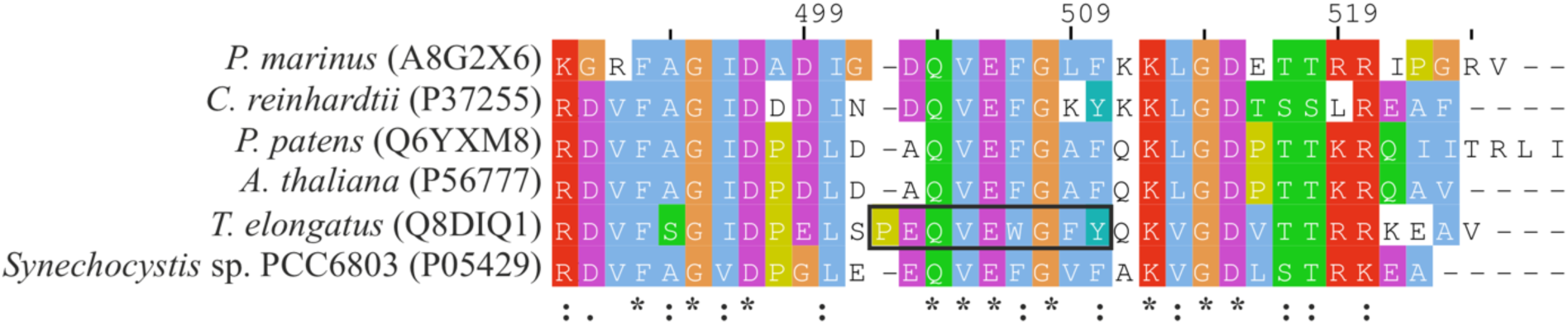
Multiple sequence alignment of the CP47 subunit from various organisms. UniProt accession codes are written in brackets. Only the C-terminal region is shown (from position 490 to the C-terminus). The black square denotates the peptide used for NMR spectroscopy. The alignment was created using the Clustal Omega web tool (Madeira et al., 2019) and visualized with Jalview (Waterhouse et al., 2009) using the ClustalX color scheme.

**Figure S5:**
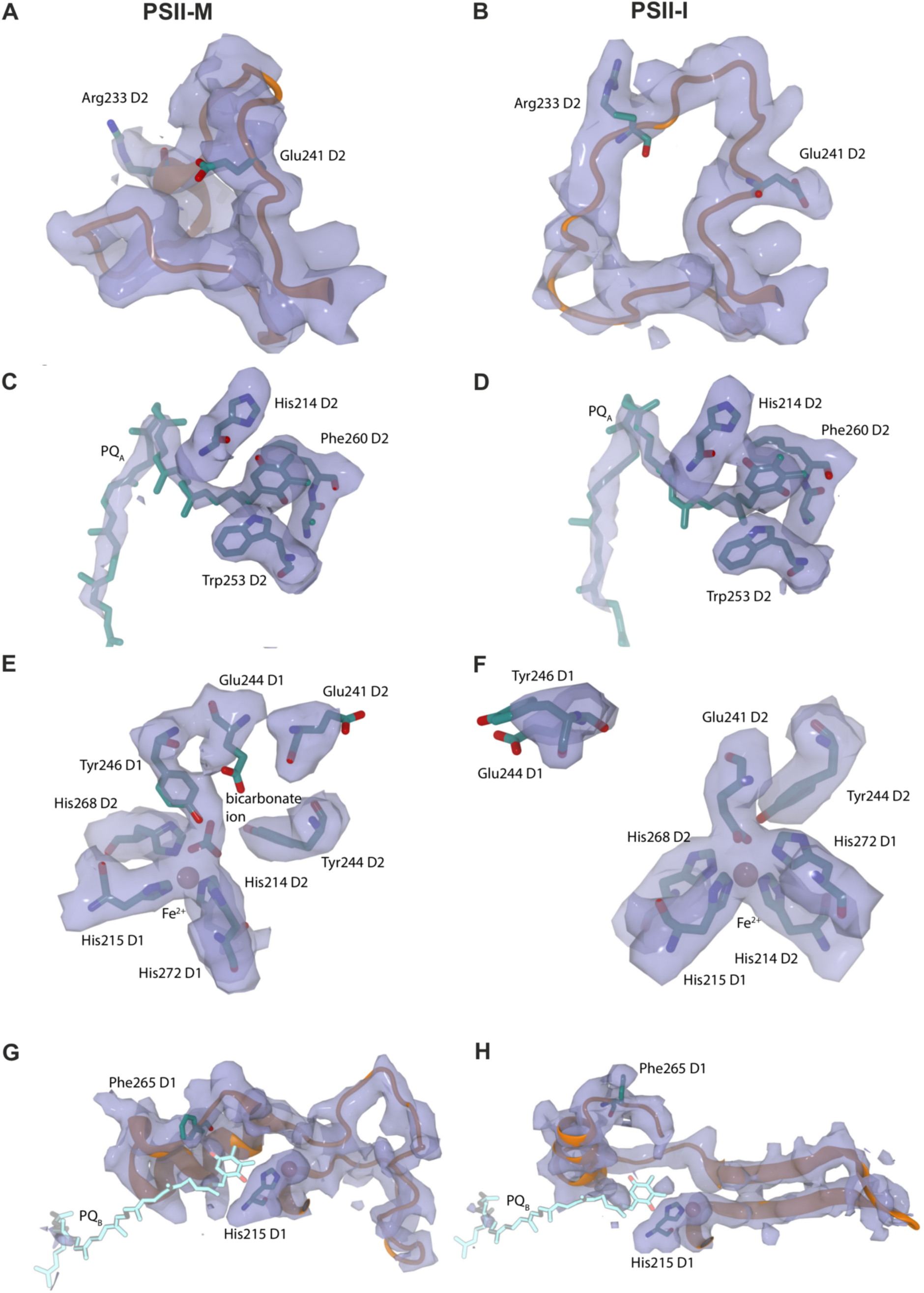
Comparison between the structural model and the density for important features at the cytoplasmic PSII side. Structural changes of the cytoplasmic D2 loop. The structural model and density (transparent blue) of the cytoplasmic D2 loop in PSII-M **(A)** and PSII-I **(B)**. Arg233 and Glu241 are highlighted to illustrate the conformational changes induced by Psb28 binding. Arg233 is one of the residues that forms the interface to Psb28 in PSII-I, and Glu241 replaces the bicarbonate ion and directly binds to the non-heme iron in PSII-I. **The Q**_**A**_ **binding pocket**. The structural model and map density (transparent blue) of the Q_A_ binding pocket in PSII-M **(C)** and PSII-I **(D)**. Key residues of D2 involved in Q_A_ binding are highlighted, including the side chain of His214 and the backbone of Phe260 that form a hydrogen bond to Q_A_, as well as the side chains of Phe260 and Trp253 that coordinate Q_A_ through Π-Π interactions. There is a clear density for Q_A_ in both maps, showing that the quinone is bound in the same conformation with and without bound Psb28. **Comparison of the non-heme Fe**^**2+**^ **coordination in PSII-M and PSII-I**. The structural model and density (transparent blue) are shown for the non-heme iron coordination sphere of PSII-M **(E)** and PSII-I **(F)**. Through binding of Psb28 in PSII-I, bicarbonate is replaced by D2 Glu241. In addition, D2 Glu244 and Tyr246 are displaced from the non-heme iron binding site. **Overview of the possible Q**_**B**_ **binding pocket**. The structural model and map density (transparent blue) of the potential Q_B_ binding pocket in PSII-M **(G)** and PSII-I **(H)**. The two residues of D1 that are involved in Q_B_ binding (Phe265 and His215 (Umena et al., 2011)) are highlighted. The Q_B_ structural model (light blue) is from the X-ray structure (PDB-ID 5MX2 (Zhang et al., 2017)) after backbone alignment with PSII-M **(G)** and PSII-I **(H)**. There is no clear density for Q_B_ in either cryo-EM map at this position. In PSII-I **(H)**, the Q_B_-coordinating residue Phe265 is clearly at a different position, showing the distortion of the Q_B_ binding site caused by Psb28 binding.

**Fig. S6:**
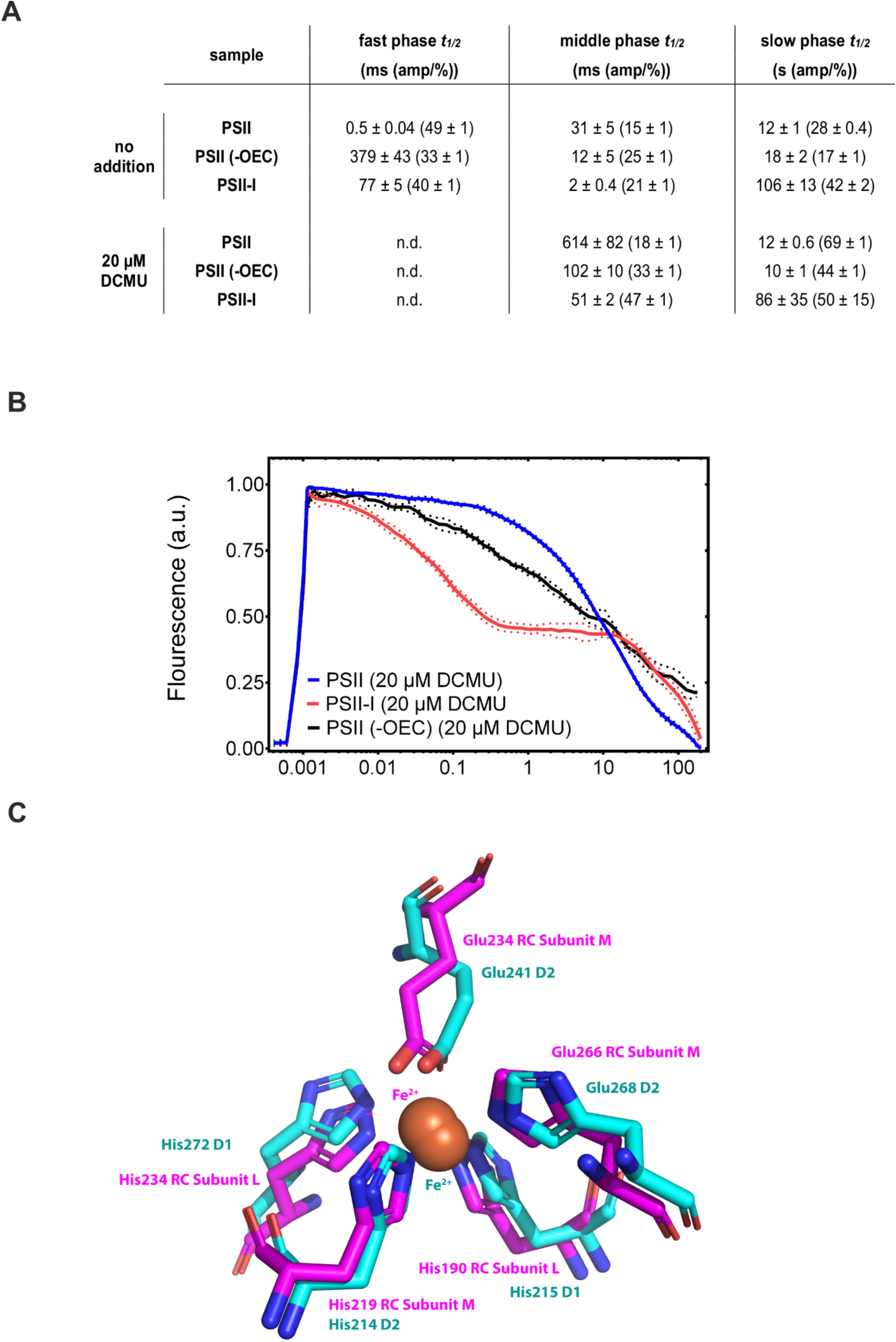
Flash-induced fluorescence decay of PSII. **(A)** Best-fit (Vass et al., 1999) decay kinetics of active PSII, inactivated PSII and PSII-I. Standard error of the calculated parameters is shown. **(B**) Fluorescence decay of PSII in the presence of 20 µM DCMU. Blue lines represent active PSII, black and red represent PSII without functional OEC and PSII-I respectively. Dotted corridors depict SD of three independent samples (*n* = 3). **(C) Comparison of the non-heme Fe**^**2+**^ **coordination in PSII-I and the non-oxygenic bacterial reaction center in *Rhodobacter sphaeroides***. Alignment of the structural model of the non-heme iron coordination sphere of PSII-I with the X-ray structure of the photosynthetic reaction center in *Rhodobacter sphaeroides* (PDB-ID 1AIJ). The structural alignment clearly reveals the structural similarities between both systems.

**Figure S7:**
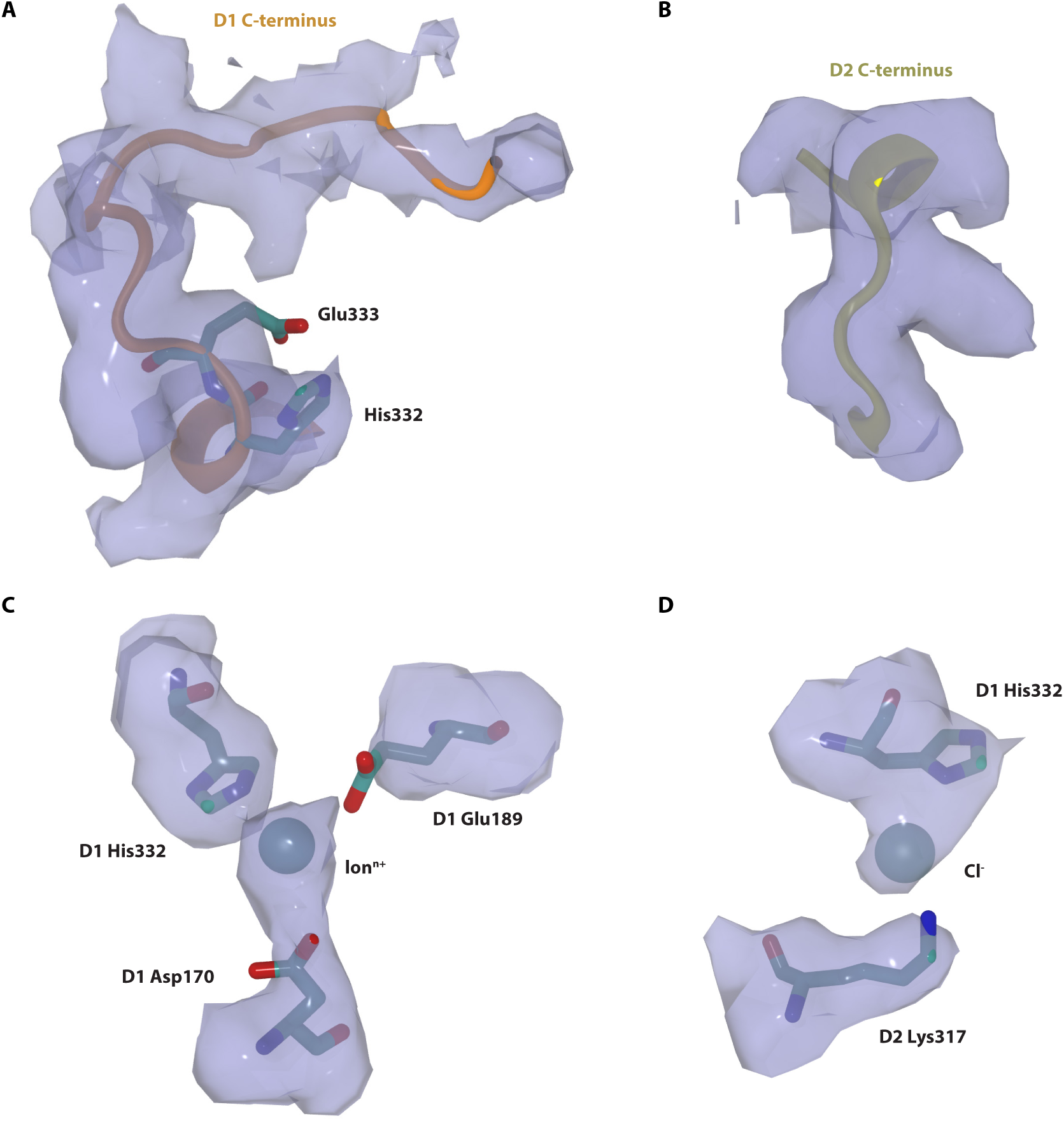
Comparison between the structural model and the density for important features at the premature OEC of PSII-I. Fit of the structural model into the map density (transparent blue) for the C-terminal tails of D1 (**A**) and D2 (**B**) of PSII-I. **C**. Ion density and its coordinating side chains in the active side of the Mn_4_CaO_5_ cluster. **(D)** Density of an ion coordinated by D1 His332 and D2 Lys317. In the available X-ray structures a Cl^-^ is assigned at this position. The resolution is high enough for a confident modeling of the C-terminal tails and ion assignment in the active site of the Mn_4_CaO_5_ cluster of PSII-I. The structural model and its fit to the density of the here shown active side of PSII-I is comparable to the one in PSII-I’ and PSII-M.

## Supplementary Tables

**Table S1:** Cryo-EM data collection, refinement, and validation statistics.

| Name | PSII-M | PSII-I | PSII-I' |
| --- | --- | --- | --- |
| Initial model used (PDB code) | 3KZI, | 3KZI<br>2YZX(Psb27)<br>3ZNP (Psb28)<br>*de novo* (Psb34) | 3KZI<br>3ZNP (Psb28)<br>*de novo* (Psb34) |
| Model resolution (Å) | 2.66 | 2.72 | 2.68 |
| FSC threshold | 0.143 | 0.143 | 0.143 |
| Map sharpening *B* factor (Å<sup>2</sup>) | -80.50 | -62.00 | -60.00 |
| Model composition |  |  |  |
| Non-hydrogen atoms | 19,806 | 21,736 | 20,925 |
| Protein atoms | 16,417 | 18,400 | 17,589 |
| Ligand atoms | 3389 | 3336 | 3336 |
| *B* factors (Å<sup>2</sup>) |  |  |  |
| Min | -69.38 | -63.43 | -64.8 |
| Mean | -111.75 | -118.19 | -119.19 |
| Max | -209.48 | -219.17 | -241.81 |
| R.m.s. deviations |  |  |  |
| Bond lengths (Å) | 0.014 | 0.015 | 0.015 |
| Bond angles (°) | 2.198 | 2.399 | 2.410 |
| Validation |  |  |  |
| MolProbity score | 1.67 | 1.98 | 1.75 |
| Clash score | 2.85 | 5.55 | 4.53 |
| EM Ringer score | 3.78 | 3.09 | 3.14 |
| Rotamer Outliers (%) | 1.89 | 2.34 | 1.62 |
| Ramachandran plot |  |  |  |
| Favored (%) | 94.241 | 94.11 | 94.53 |
| Allowed (%) | 5.45 | 5.62 | 5.19 |
| Outliers (%) | 0.15 | 0.26 | 0.28 |

**Table S2:**
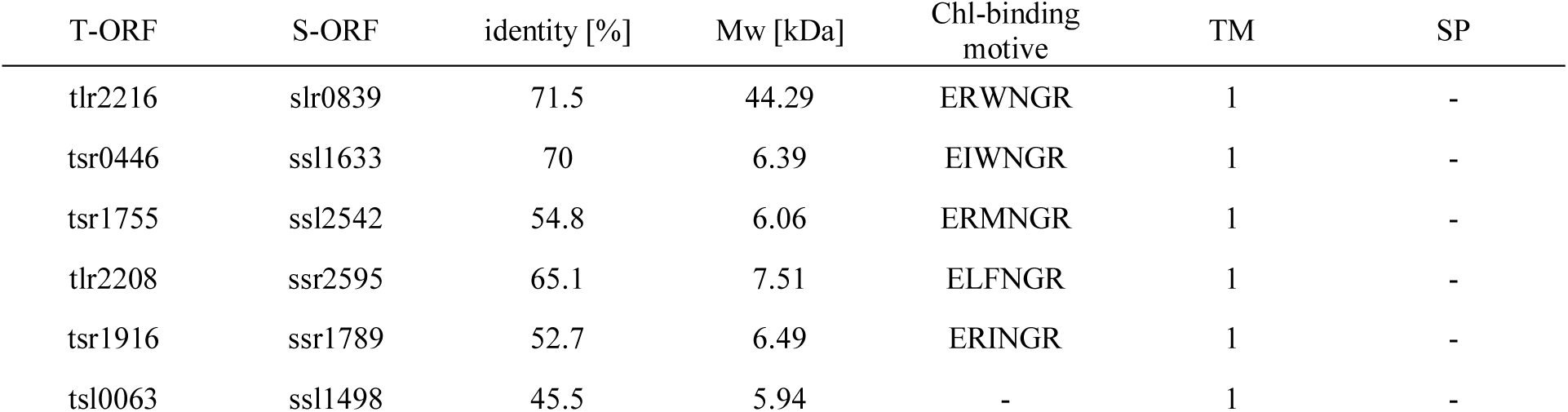
High-light inducible proteins in *T. elongatus*. Listed are the open reading frames (ORF) in *T. elongatus* (T) and in *Synechocystis sp*. PCC6803 (S), sequence identity and molecular weight (Mw) of the translated proteins, sequence of the chlorophyll binding motive, number of trans-membrane helixes (TM) and whether the sequence possesses a signal peptide for import into the thylakoid lumen (SP).

## Supplementary Movies

**Movie 1**.

Rigid body rotation and attachment of the CP43 antenna after Psb28 dissociation (morph of PSII-I and PSII-M). Accompanies Fig. 3A, Fig. 4A and B.

**Movie 2**

Q_B_ binding site maturation (morph of PSII-I and PSII-M). Reorientation of D1 Phe265 after Psb28 dissociation. The structural model of Q_B_ was added from the X-ray structure (PDB-ID 3WU2) after backbone alignment with PSII-M. Accompanies Fig. 4E and F.

**Movie 3**

Reorientation of D2 Glu241 and bicarbonate binding after Psb28 dissociation (morph of PSII-I and PSII-M). Accompanies Fig. 5A-D.

**Movie 3**

Maturation of the oxygen evolving cluster (morph of PSII-I and mature PSII (PDB-ID 3WU2)). Reorientation of the D1 C-terminus (colored in *light blue*, His332, Glu333, His337, Asp342, Ala344) and additional residues involved in binding of the potential Cl^-^ (D2 Lys317) as well as of the potential Mn^2+^ ion (D1 Asp170 and Glu189). Accompanies Fig. 6A and Fig. 7.

## Methods

### Cultivation of *Thermosynechococcus elongatus* BP-1

Cell growth and thylakoid membrane preparation were performed as described previously (Kuhl et al., 2000). In brief, *T. elongatus* mutant strains (ΔpsbJ psbC-TS and psb34-TS) were grown in BG-11 liquid medium inside a 25-liter foil fermenter (Bioengineering) at 45°C, 5% (v/v) CO_2_-enriched air bubbling and 50-200 µmol photons m^−2^ s^−1^ white light illumination (depending on the cell density). Cells were harvested at an OD_680_ of ∼ 2 after 5-6 days of cultivation and concentrated to ∼ 0.5 l using an Amicon DC10 LA hollow fiber system, pelleted (3500 rcf, 45 min and 25 °C) and resuspended in 150 ml of Buffer D (100 mM Tris-HCL, pH 7.5, 10 mM MgCl_2_, 10 mM CaCl_2_, 500 mM mannitol and 20% (w/v) glycerol). The harvested cells were flash frozen in liquid nitrogen and stored at –80 °C until further use.

### Preparation of *T. elongatus* mutant strains

*Thermosynechococcus elongatus* ΔpsbJ psbC-TS was generated based on the previously described strain *T. elongatus* ΔpsbJ (Nowaczyk et al., 2012) that was transformed with the plasmid pCP43-TS. The plasmid is based on pCP34-10His (Nowaczyk et al., 2006). The His-tag sequence was exchanged with TwinStrep-tag by PCR using the primers CP43TS_rev (5’CCCGATATCTTACTTCTCAAATTGCGGATGAGACCACGCAGAACCACCAGAACCACCGCCGCTGCCGCCGCCTTTTTCGAACTGCGGGTGGCTCC 3’) and NTCP43 (5’ TGCTCTA GAATGAAAACTTTGTCTTCCCAGA 3’). The resulting PCR product was ligated back into an empty pCP34-10His backbone using XbaI and EcoRV restriction endonucleases. *T. elongatus* BP-1 cells were transformed as described previously (Iwai et al., 2004). Mutant colonies were selected by frequent re-plating onto agar plates with increasing antibiotic concentrations, stopping at 8 µg/ml of chloramphenicol and 80 µg/ml of kanamycin. Complete segregation of the mutant was confirmed by PCR with the primers CTCP43DS (5’ CCGCTCGAGACCATCCAAGCTTGGCAGCA 3’) and NTCP43 (5’ TGCTCTAGAAT GAAAACTTTGTC TTCCCAGA 3’). *T. elongatus* psb34-TS was generated by transformation with the plasmid pPsb34-TS. The plasmid DNA was obtained from TwistBioscience. It consisted of psb34 (tsl0063) with a C-terminal TwinStrep-tag and a kanamycin resistance cassette, flanked by tsl0063-upstream and downstream regions (900 bp each). *T. elongatus* BP-1 cells were transformed (Iwai et al., 2004) and mutant selection took place (Nowaczyk et al., 2006). Complete segregation of the mutant was verified by PCR. The primers used were tsl0063-up-for (5’ CATATGGTCTCGCAATTATTTGCCCATGC 3’) and tsl0063-down-rev (5’ GGTACCCCGACACAGTTGATCACCGC 3’).

### Purification of photosystem II assembly intermediates

Thawed cells were diluted in 100 ml of Buffer A (100 mM Tris-HCL, pH 7.5, 10 mM MgCl_2_ and 10 mM CaCl_2_) and pelleted again (21 000 rcf, 20 min and 4°C). The pellet was resuspended in 100 ml of Buffer A with 0.2% (w/v) lysozyme and dark incubated for 75-90 min at 37 °C. This was followed by cell disruption by Parr bomb (Parr Instruments Company) and pelleting (21 000 rcf, 20 min and 4°C). All following steps were performed under green illumination. The pellet was resuspended in 150 ml of Buffer A and pelleted again (21 000 rcf, 20 min and 4°C). This step was repeated three times, with the last resuspension in 80 ml of Buffer B (100 mM Tris-HCL, pH 7.5, 10 mM MgCl_2_, 10 mM CaCl_2_ and 500 mM mannitol). The isolated thylakoids were flash frozen in liquid nitrogen and stored at −80 °C.

Strep-Tactin-affinity purification of PsbC-TS and Psb34-TS assembly intermediates were performed under green illumination. Membrane protein extraction was performed as described previously (Kuhl et al., 2000), with certain adaptations. Thylakoid membranes were supplemented with 0.05% (w/v) n-Dodecyl β-maltoside (DDM) (Glycon) and pelleted (21000 rcf, 20 min and 4°C). The sample was resuspended in extraction buffer (100 mM Tris-HCL, pH 7.5, 10 mM MgCl_2_, 10 mM CaCl_2_, 1.2% (w/v) DDM, 0.5% (w/v) sodium-cholate and 0.01% (w/v) DNase) to a final chlorophyll concentration of 1 mg/ml and incubated for 30 min at 20 °C. The solubilized membrane proteins were ultra-centrifugated (140000 rcf, 60 min and 4 °C) and NaCl was added to the supernatant to a final concentration of 300 mM.

The supernatant was filtered through a 0.45 µm filter and applied to a 5 ml Strep-Tactin Superflow HC column (IBA Lifesciences), equilibrated in Buffer W (100 mM Tris-HCL, pH 7.5, 10 mM MgCl_2_, 10 mM CaCl_2_, 500 mM mannitol, 300 mM NaCl and 0.03% (w/v) DDM) at a flowrate of 3 ml/min. The column was washed with Buffer W until a stable baseline (A280) was reached. Strep-tagged protein complexes were eluted by an isocratic elution with Buffer E (100 mM Tris-HCL, pH 7.5, 10 mM MgCl_2_, 10 mM CaCl_2_, 500 mM mannitol, 300 mM NaCl 2.5 mM desthiobiotin and 0.03% (w/v) DDM). The captured fractions were equilibrated in Buffer F (20 mM MES, pH 6.5, 10 mM MgCl_2_, 10 mM CaCl_2_, 500 mM mannitol and 0.03% (w/v) DDM) with a spin concentrator (Amicon, Ultra – 15, 100000 NMWL), flash-frozen in liquid nitrogen and stored at −80 °C until analysis.

PsbC-TS containing assembly intermediates were further separated by ion exchange chromatography (IEC). Captured elution fraction from the Strep-Tactin-affinity purification were loaded onto a anion exchange column (UNO Q-6, Biorad) with a flowrate of 4 ml/min, pre-equilibrated in Buffer F. Protein complexes were eluted by a liner gradient of MgSO_4_ (0-150 mM) using Buffer G (20 mM MES, pH 6.5, 10 mM MgCl_2_, 10 mM CaCl_2_, 500 mM mannitol, 150 mM MgSO_4_ and 0.03% (w/v) DDM). Fractions containing PSII assembly intermediates were collected, concentrated to 100 – 10 µM reaction centers, using a spin concentrator (Amicon, Ultra – 15, 100 000 NMWL), aliquoted, flash frozen in liquid nitrogen and stored at −80 °C until further analysis.

### Protein Expression and Purification of Psb28

The Psb28 expression plasmid was constructed by first amplifying psb28 from *T. elongatus* genomic DNA, using primers TeloPsb28for (5’ GGAATTCCATATGGGTGCAATGGCAGAAATTC 3’) and TeloPsb28rev (5’ CGAATTCCCCGGGAGAGTTCTCAGACTTCTG 3’). Next, the amplified DNA was cloned into pIVEX2.3d using *Nde*I/*Sma*I to obtain pIVEXPsb28His. Expression and purification of ^15^N-labelled Psb28 was carried out as described previously (Cormann et al., 2014) with certain adaptations. Overnight starter cultures were grown on agar plates at 37 °C, supplemented with 1 % (w/v) glucose and 100 µg/ml ampicillin. The cell material was then resuspended in 2 ml of M9 media (Studier, 2005), and this was used to inoculate 500 ml of M9 media with ^15^NH_4_Cl as the only nitrogen source. Cultures were incubated at 37 °C under vigorous shaking and at an OD_600_ of 0.6, and expression was induced with addition of isopropylthiogalactoside to a final concentration of 0.5 mM. After overnight incubation the ^15^N-labbled Psb28 was isolated and purified as described for his-tagged CyanoP (Cormann et al., 2014). The purity and integrity of the protein samples were checked by SDS-PAGE (data not shown).

### Polyacrylamide Gel Electrophoresis

Blue-native PAGE (Neff and Dencher, 1999) was used to assess the oligomeric state of the isolated PSII assembly intermediates. Separation of protein complexes was carried out across a linear gradient of polyacrylamide (acrylamide-bisacrylamide, 32:1) from 3.2 to 16% (w/v) in the separating gel. This was overlaid with a 3% (w/v) polyacrylamide (acrylamide-bisacrylamide, 32:1) sample gel. The gels were loaded with 40 µg of protein per lane. Electrophoresis was performed at 4 °C in a Mini-PROTEAN Tetra System (BioRad) at 100 V for 30 min with Blue Cathode Buffer (15 mM BisTris-HCl, pH 7.0, 50 mM Tricine and 0.002% (w/v) Coomassie Brilliant Blue 250) and at 170 V for an additional 90 min with Cathode Buffer (15 mM BisTris-HCl, pH 7.0 and 50 mM Tricine). The anode buffer was composed of 50 mM BisTris-HCl at pH 7.

Subunit composition was investigated by SDS-PAGE (Schagger and von Jagow, 1987). Separation of polypeptide chains took place on a 19% (w/v) polyacrylamide gel (acrylamide-bisacrylamide, 37.5:1), containing 9 M urea and 4% (w/v) glycerol. The gel was loaded with 40 µg of denatured protein complex per lane. The gels ran at 4 °C in a Mini-PROTEAN Tetra System (BioRad) at 35 mA per gel for 60 min. Fixation and visualization of polypeptide chains was performed with Coomassie Staining Solution (45% (w/v) isopropanol, 10% (w/v) acetate and 0.2% (w/v) Coomassie Brilliant Blue 250).

### Mass spectrometry analysis

PSII-I complexes were purified and desalted using Isolute C18 SPE cartridges (Biotage, Sweden). The columns were first washed and equilibrated, the sample diluted in 0,1% trifluoroacetic acid (TFA) and loaded onto the column. After washing with 2 ml 0.1% TFA, the proteins were eluted with 500 µl 80% acetonitrile (ACN), 20% water. The organic fraction was lyophilized in a vacuum concentrator (Eppendorf, Germany), reconstituted in 0.1% TFA and mixed in a 1:1 ratio with HCCA matrix solution (HCCA (alpha-cyano-4-hydroxycinnamic acid) saturated in 50% ACN, 50% water and supplemented with 0.1% TFA). Subsequently, 1 µl aliquots of the mixture were deposited on a ground steel MALDI target and allowed to dry and crystallize at ambient conditions.

MS and MS/MS spectra were acquired on a prototype rapifleX MALDI-TOF/TOF (Bruker Daltonics, Germany) in positive ion mode. The Compass 2.0 (Bruker Daltonics, Germany) software suite was used for spectra acquisition and processing (baseline subtraction, smoothing, peak picking), a local Mascot server (version 2.3, Matrixscience, UK) was used for database searches against the *T. elongatus* proteome (UniProt, retrieved 4/2019) and BioTools 3.2 (Bruker Daltonics) was used for manual spectrum interpretation, de novo sequencing and peak annotation.

### Flash-induced fluorescence decay measurements

Flash-induced fluorescence decay was measured on a FL3500 Dual-Modulation Kinetic Fluorometer (PSI Photon Systems Instruments). Reaction centers were exited with 625 nm LEDs for both actinic (50 µs) and measuring flashes. The first data point was collected 80 µs after the actinic flash. Data points were collected from 80 µs to 50 or 200 s after the actinic flash for measurements with whole cells and isolated PSII, respectively. 10 data points were collected per logarithmic decade. Assays were performed at room temperature in the presence and absence of 20, 100, 200 or 400 µM 3-(3,4-dichlorophenyl)-1,1-dimethylurea (DCMU) with 5 min of dark incubation prior to measurement. Assays with isolated PSII complexes were carried out with 200 nM reaction centers in activity buffer (100 mM KCl, 20 mM MES-KOH, pH 6.5, 10 mM MgCl_2_, 10 mM CaCl_2_ and 0.03% (w/v) DDM).

### NMR Spectroscopy

Typically, NMR samples contained up to 1 mM of protein in 20 mM Tris/HCl pH 8, 10%D_2_O, 0.02% NaN_3_, and DSS. NMR spectra were acquired at 298 K on Bruker DRX 600 and AVANCE III HD 700 spectrometers. Backbone assignments for the free form of Psb28 were obtained from three-dimensional HNCA (Grzesiek and Bax, 1992; Schleucher et al., 1993), and CBCA(CO)NH (Grzesiek and Bax, 1993) spectra. Side-chain assignments were obtained from three-dimensional ^1^H-^15^N HNHA (Vuister and Bax, 1993, 1994), ^1^H-^13^C-HCCH-TOCSY (Kay et al., 1993), ^1^H-^15^N-HSQC-TOCSY (Sattler et al., 1992), ^1^H-^15^N-HSQC-NOESY (Davis, 1995), ^1^H-^13^C-HSQC-NOESY (Davis et al., 1992), and aromatic ^1^H-^13^C-HSQC-NOESY spectra. Spectra were processed with NMRPipe (Delaglio et al., 1995) and analysed with CcpNmr Analysis (Vranken et al., 2005). NMR experiments for the complex form of Psb28 and the C-terminal peptide of CP47 were carried out on a Bruker AVANCE III HD 700 spectrometer at 298 K in 20 mM Tris/HCl pH8, 10%D_2_O, 0.02% NaN_3_, and DSS. For the backbone assignments of the complex, a 1 mM sample of [U-^15^N-^13^C]-enriched Psb28 was mixed with a three-fold excess of peptide. Three-dimensional (3D) HNCA, HNCO (Grzesiek and Bax, 1992; Schleucher et al., 1993), HN(CO)CACB (Yamazaki et al., 1994), HNCACB (Muhandiram et al., 1993; Wittekind and Mueller, 1993), and HN(CA)CO (Clubb et al., 1992) were recorded with 16 scans and 25% non-uniform sampling (NUS). (H)CC(CO)NH and H(CCCO)NH spectra (Logan et al., 1993; Lyons and Montelione, 1993) were recorded with 64 scans and 25% NUS. HNHA and ^1^H^15^N-HSQC-NOESY spectra were recorded with 32 scans and 25% NUS as well as traditional acquisition schemes, respectively. The mixing time for NOESY spectra was set to 120 ms. In addition, heteronuclear two-dimensional ^15^N{^1^H}-NOE data were recorded in order to extract pico-to nanosecond dynamics (Barbato et al., 1992; Ross et al., 1993). The titrations were carried out by adding increasing amounts of a peptide stock solution to the NMR sample containing 0.138 mM of protein and two-dimensional ^1^H^15^N-HSQC spectra (Mori et al., 1995) were recorded after thorough mixing of the Psb28-CP47 carboxyterminal peptide solution. Spectra were processed with NMRPipe (Delaglio et al., 1995) and Psb28 ligand affinity calculations based on two-dimensional lineshape analysis were carried out using the TITAN software package (Waudby et al., 2016).

### Synthetic Peptide

The carboxy-terminal peptide from residues 480-499 of CP47, which comprises the sequence SGIDPELSPEQVEWGFYQKV and includes an acetylated amino-terminus, was purchased from JPT Peptide Technologies GmbH, Germany. Peptide stock solutions of at least 6.07 mM for titration experiments were prepared by dissolving the peptide in 20 mM Tris/HCl pH8.

### Removal of the PSII oxygen evolving cluster

PSII without functional oxygen evolving cluster (OEC) was prepared by isolating PSII as described by Grasse et al. (Grasse et al., 2011), followed by removal of the extrinsic subunits according to Shen and Inoue (Shen and Inoue, 1993), with modifications. PSII was applied to a size exclusion column (Superdex 75 10/300 GL, GE Healthcare) pre-equilibrated in CaCl_2_ buffer (10 mM MgCl_2_, 20 mM MES-NaOH, pH 6.5, 1 M CaCl_2_, 0.03% (w/v) DDM). PSII particles lacking the extrinsic subunits and the Mn_4_CaO_5_ cluster, which were eluted in the void volume, were collected and the buffer was exchanged to activity buffer (100 mM KCl, 20 mM MES-KOH, pH 6.5, 10 mM MgCl_2_, 10 mM CaCl_2_ and 0.03% (w/v) DDM) using a spin concentrator (Amicon, Ultra – 15, 100 000 NMWL).

### Detection of singlet oxygen by the room temperature EPR spectroscopy

Singlet oxygen was trapped using the water-soluble spin-probe 2,2,6,6-tetramethyl-4-piperidone (TEMPD) hydrochloride (Hideg et al., 2011) and measured with ESR300 (Bruker Biospin, Rheinstetten, Germany). Samples (30 µg chl ml−1) were illuminated for 1 min with red light (RG 630) at 450 µmol quanta m^-2^s^-1^ in 0.5 M mannitol, 10 mM CaCl_2_, 10 mM MgCl_2_, 20 mM MES at pH 6.5. Spectra were recorded using a flat cell containing 200 µl sample. The microwave power was 9.77 GHz and 14.07 mW with a modulation frequency of 86 kHz and amplitude of 1.0 G. Each spectrum is an average of 8 scans, each with a sweep time of 10.5 s.

### Cryo-electron microscopy

For cryo-EM sample preparation, 4.5 µl of purified protein complexes were applied to glow discharged Quantifoil 2/1 grids, blotted for 3.5 s with force 4 in a Vitrobot Mark III (Thermo Fisher) at 100% humidity and 4°C, then plunge frozen in liquid ethane, cooled by liquid nitrogen. Cryo-EM data was acquired with a FEI Titan Krios transmission electron microscope using the SerialEM software (Mastronarde, 2005). Movie frames were recorded at a nominal magnification of 22,500x using a K3 direct electron detector (Gatan), The total electron dose of ∼55 electrons per Å2 was distributed over 30 frames at a calibrated physical pixel size of 1.09 Å. Micrographs were recorded in a defocus range of −0.5 to −3.0 µm.

### Image processing, classification and refinement

Cryo-EM micrographs were processed on the fly using the Focus software package (Biyani et al., 2017) if they passed the selection criteria (iciness < 1.05, drift 0.4 Å < x < 70 Å, defocus 0.5 um < x < 5.5 um, estimated CTF resolution < 6 Å). Micrograph frames were aligned using MotionCor2 (Zheng et al., 2017) and the contrast transfer function (CTF) for aligned frames was determined using Gctf (Zhang, 2016). Using Gautomatch (http://www.mrc-lmb.cam.ac.uk/kzhang/) 693,297 particles were picked template-free on 824 acquired micrographs. Particles were extracted with a pixel box size of 260 using RELION 3.1 (Scheres, 2012) and imported into Cryosparc 2.3 (Punjani et al., 2017). After reference-free 2D classification, 675,123 particles were used for ab initio construction of initial models and subjected to multiple rounds of 3D classification to obtain models with and without Psb28 density. Non-uniform refinement in Cryosparc resulted in models with an estimated resolution of ∼3.2 Å. Particles belonging to 3D classes with and without Psb28 (150,090 and 166,411 particles, respectively) were reextracted in RELION with a pixel box size of 256 and subjected to several rounds of CTF-refinement (estimation of anisotropic magnification, fit of per-micrograph defocus and astigmatism and beam tilt estimation) and Bayesian polishing (Scheres, 2014). Both classes were refined using the previously generated starting models. 3D classification without further alignment using a mask around the Psb27 region separated particles in the Psb28-containing class into distinct classes with and without Psb27 (57,862 and 91,473 particles, respectively). Final refinement of each of the three classes (with Psb27 and Psb28 (PSII-I), with Psb28 but without Psb27 (PSII-I’), and without Psb27 and Psb28 (PSII-M)) resulted in models with global resolutions of 2.94 Å, 2.76 Å and 2.82 Å, respectively (Gold standard FSC analysis of two independent half-sets at the 0.143 cutoff). Local-resolution and 3D-FSC plots (Extended Data Fig. 2) were calculated using RELION and the “Remote 3DFSC Processing Server” web interface (Tan et al., 2017), respectively.

### Atomic model construction

The 3.6 Å resolution X-ray structure of monomeric PSII from *T. elongatus* with PDB-ID 3KZI (Broser et al., 2010) was used as initial structural model that was docked as rigid body using Chimera (Pettersen et al., 2004) into the obtained cryo EM densities for PSII-M and PSII-I. The cofactors that had no corresponding density were removed. The subunit PsbJ was also removed, as it was deleted in the experimental design. By highlighting the still unoccupied parts of the PSII-I density map, we identified densities that lead to the structures of Psb27, Psb28, and Psb34.

The 2.4 Å resolution X-ray structures of isolated Psb28 from *T. elongatus* with PDB-ID 3ZPN (Bialek et al., 2013) and the 1.6 Å resolution X-ray structure of isolated Psb27 from *T. elongatus* with PDB-ID 2Y6X (Michoux et al., 2012) were docked as rigid bodies into the unoccupied densities. The 1.6 Å resolution X-ray structure of CyanoQ from *T. elongatus* with PDB-ID 3ZSU (Michoux et al., 2014) does not fit into the density and was therefore not modeled.

As there was no experimentally resolved structural model of Psb34 available, we first used the sequence with UniProt-ID Q8DMP8 to predict structures using the webserver SWISS Model (Schwede et al., 2003) and LOMETS (Wu and Zhang, 2007). We also predicted the secondary structure through the meta server Bioinformatics Toolkit (Zimmermann et al., 2018) and CCTOP (Dobson et al., 2015). The results of the secondary structure prediction are summarized in Table S4. Combining these predictions together with the unassigned cryo-EM density, we used COOT (Emsley et al., 2010) to build an initial model of Psb34 that has one α-helix from amino acid number 28 to 55.

### Model Refinement

The initial model of the complex described above was refined in real space against the cryo-EM density of PSII-I, and structural clashes were removed using molecular dynamics flexible fitting (MDFF) (Trabuco et al., 2009). MDFF simulations were prepared in VMD 1.9.4a35 (Humphrey et al., 1996) using QwikMD (Ribeiro et al., 2016) and the MDFF plugin. The simulations were carried out with NAMD 2.13 (Phillips et al., 2005) employing the CHARMM36 force field. Secondary structure, cis peptide and chirality restraints where employed during 800 steps of minimization followed by a 40 ps MDFF simulation at 300K. Due to the employed restraints, only conformational changes of side chains and subunit movements compared to the initial structure are identified during the initial MDFF run. We checked the fit to density of the structure by calculating cross-correlation values of the backbone atoms. For PSII-I, we identified residues 217 to 269 from PsbA and residues 467 to 499 from PsbB and PsbZ as main regions where the structural model was not yet in accordance with the density after the initial MDFF run. For these three regions, we employed an iterative combination of MDFF with Rosetta (Leaver-Fay et al., 2011; Lindert and McCammon, 2015). Here, we used the optimized strategy as described for model construction of the 26S proteasome (Guo et al., 2018; Wehmer et al., 2017).

To obtain an atomic model that fit the PSII-M density, we used the initial model based on 3KZI described above, but without PsbJ, Psb27, Psb28, and Psb34. After the initial MDFF run, the cross-correlation check did not reveal any regions with significant deviation between model and density. Therefore, no further refinement was necessary. This fast convergence reflects that there are no crucial differences between the PSII-M model and the X-ray structure 3KZI.

To obtain the atomic model that fit the PSII-I’ density, we used the final PSII-I model without Psb27 for MDFF. After the initial MDFF run, the cross-correlation check did not reveal any regions with significant deviation between model and density. This fast convergence reflects that there are no crucial differences between the PSII-I and PSII-I’ models, except for the presence of the Psb27 subunit.

Last, the PSII-M, PSII-I, and PSII-I’ models were used to initiate one final round of real-space refinement in Phenix (Liebschner et al., 2019).

## Acknowledgments

We thank C. König, M. Völkel, and R. Oworah-Nkruma for excellent technical assistance, Kristin Becker for cloning of the pIVEXPsb28His plasmid, Bibi Erjavec for preparation of the scheme in Fig. 1 and Nicholas Cox for helpful discussion. J.M.S. is grateful to E. Conti for scientific independence and great mentorship and to J. M. Plitzko and W. Baumeister for access to the cryo-EM infrastructure and early career support. M.M.N. is grateful to his mentor M. Rögner for generous support.

## Author contributions

B.D.E., T.R., J.M.S. and M.M.N. conceived the research, prepared the figures, and wrote the manuscript with the contributions of all other authors. M.M.N. coordinated the activities. Preparation of mutants, PSII isolation and biochemical analysis were performed by J.Z., M.M, P.L. and M.M.N. Mass spectrometry analysis was done by J.M.-C. and J.D.L. J.M.S., S.B. and B.D.E. performed the cryo-EM analysis and T.R. built the structural model with the help of S.K.S., A.C. and E.T. Fluorescence spectroscopy was carried out by J.Z. and M.M.N. EPR experiments were conducted by A.K.-L. NMR experiments were conducted and analyzed by O.A. and R.S.. All authors approved the final version of the manuscript.

## Data availability

The cryo-EM density maps will be deposited in the Electron Microscopy Data Bank, the atomic models of the cryo-EM structures in the worldwide Protein Data Bank (wwPDB) and the NMR assignments for Psb28 in the Biological Magnetic Resonance Bank (BMRB), respectively.

